# Mitochondrial amino acid export preserves mitochondrial volume at the egg-to-embryo transition

**DOI:** 10.64898/2026.10.08.757432

**Authors:** Melissa C. Pamula, Lauren Clamon, Samuel Block, Suzanne Savage, Christoph Gaebelein, Ashid Amarsanaa, Margaret E Bisher, David Mankus, Abigail KR Lytton-Jean, Nora Kory, Ruth Lehmann

## Abstract

In animals, the transition from egg to embryo marks a rapid shift in cellular physiology, yet how mitochondria adapt to support this conversion remains poorly understood. We performed a targeted RNAi screen of conserved, nuclear-encoded mitochondrial proteins in *Drosophila melanogaster* to identify maternal effects. We identified distinct phenotypic classes affecting mitochondrial distribution or morphology in early embryos. We focused on Sideroflexin-1/3 (Sfxn1-3), a conserved mitochondrial inner membrane transporter, because it uniquely affects embryonic mitochondrial morphology. Maternal loss of Sfxn1-3 causes striking mitochondrial enlargement in embryos but not in oocytes, placing the trigger at the egg-to-embryo transition. Combining mitochondrial immunopurification with LC/MS-based metabolomic profiling, we show that amino acids are markedly enriched in Sfxn1-3-deficient mitochondria in both oocytes and embryos. This accumulation developmentally uncouples the metabolic and morphological phenotypes. By selectively disrupting discrete steps of egg activation, including translational activation, cell cycle re-entry, and fertilization, we demonstrate that mitochondrial swelling is independent of each, prompted instead by egg hydration. Beyond its reported role in proliferating cells as a serine importer, our data support a role for germline Sfxn1-3 in regulating inner mitochondrial membrane permeability to osmotically active amino acids. This activity is crucial for preserving organelle volume during the osmotic challenge of oocyte hydration. When this export fails, amino acid accumulation amplifies osmotic stress and drives rapid mitochondrial swelling. Together, these findings identify mitochondrial inner membrane permeability to amino acids as a key regulatory step that couples pre-established metabolite compartmentalization to osmotic adaptation at egg activation, and reveal how mitochondrial metabolic integrity and volume control are coordinated at this critical developmental transition.

## Introduction

In sexually reproducing species, mitochondria are transmitted from one generation to the next exclusively through the maternal gamete. In the early stages of oogenesis, purifying mitochondrial selection serves as a quality-control mechanism unique to germ cells, counteracting the otherwise irreversible decay of the mitochondrial genome across generations (Z. Chen & Xu, 2026; Lieber et al., 2019). The mature egg provides the entire mitochondrial complement of the future embryo. Oocytes suppress electron transport chain activity, a mechanism hypothesized to help preserve the tissue by minimizing oxidative stress, and this suppression is relieved in the embryo where energetic demands rapidly increase (Berg et al., 2024; Dumollard et al., 2007). While global metabolic remodeling has been studied during the maternal-to-zygotic transition (Chugunova et al., 2026; Pérez-Mojica et al., 2025), less is known about how mitochondria themselves adapt to these changes.

Beyond their metabolic roles, mitochondria must maintain their structural integrity as osmotic conditions in the cell change. Matrix volume is not a passive property of the organelle: solute accumulation in the matrix draws water inward, and the resulting swelling can disorganize cristae, uncouple respiration, and, in extreme cases, rupture the outer membrane (Bernardi et al., 2023; Sesso et al., 2012). Cells therefore rely on regulated transport of matrix ions and metabolites to prevent pathological swelling, particularly during rapid shifts in cytoplasmic osmolarity. The egg-to-embryo transition is an extreme, naturally occurring example of such a shift: the mature oocyte, which is dehydrated during late oogenesis, undergoes rapid hydration prior to fertilization, causing cytoplasmic osmolarity to abruptly drop (Horner & Wolfner, 2008a; Mahowald et al., 1983; York-Andersen et al., 2021). How organelles, and mitochondria in particular, maintain volume homeostasis during this and other rapid physiological transitions remains unknown.

We chose to study mitochondrial volume control at the egg-to-embryo transition in *Drosophila melanogaster*, where oocyte maturation, egg activation, and fertilization are discrete, sequential events that can each be genetically perturbed and analyzed. During mid- to late-stage oogenesis, meiotically arrested oocytes progressively dehydrate. Egg activation begins when the mature oocyte enters the oviduct and encounters hypotonic fluid. This triggers oocyte swelling mediated by aquaporins and DEG/ENaC channels, followed by constriction that mechanically initiates egg activation (Heifetz et al., 2001; Horner & Wolfner, 2008b, 2008a; Page & Orr-Weaver, 1997). A transient calcium wave sweeps through the egg, licensing a coordinated set of activities, including meiotic completion, Pan Gu (Png) kinase-dependent translational activation of maternal mRNAs, and Cdk1-dependent cell cycle re-entry (Avilés-Pagán & Orr-Weaver, 2018; Hara et al., 2018; Hu & Wolfner, 2019; Kaneuchi et al., 2015; York-Andersen et al., 2021). These events proceed independently of fertilization, which involves nuclear reprogramming and initiates embryonic development. Because hydration, activation, and fertilization can each be manipulated independently, this system allows for the dissection of how mitochondria are affected by and respond to specific physiological triggers.

We identified Sideroflexin 1/3 (Sfxn) as a determinant of mitochondrial volume control during the egg-to-embryo transition in *Drosophila melanogaster*. We show that Sfxn buffers mitochondria against osmotic stress by preventing the accumulation of neutral amino acids in the mitochondrial matrix. High-resolution light and electron microscopy reveal that Sfxn-deficient mitochondria are isotropically and massively enlarged in early embryos, but morphologically normal immediately before egg hydration, placing mitochondrial swelling at egg activation. Combining mitochondrial immunopurification with metabolomics, we find that neutral amino acids accumulate inside Sfxn-depleted mitochondria, a signature inconsistent with a simple import defect. Neither the swelling nor the amino acid accumulation phenotype can be attributed to impaired one-carbon metabolism, which yields a distinct metabolic and phenotypic signature. Similarly, respiratory chain complex assembly and function are not impaired. These results support a model in which maternally supplied Sfxn prevents free (e.g., osmotically active) amino acids from accumulating in the matrix during oogenesis, thereby preventing a matrix solute burden from becoming pathological as cytoplasmic osmotic pressure drops during egg hydration.

Together, these findings identify mitochondrial inner-membrane amino acid permeability as a control point that couples metabolite compartmentalization to organelle volume homeostasis at a critical developmental transition.

## Results

### Maternal mitochondrial proteins shape mitochondrial pattern and morphology in the embryo

To assess the maternal contribution to mitochondrial form and function in the embryo, we performed an *in vivo* RNAi screen targeting maternally expressed mitochondrial genes in *Drosophila melanogaster*. The mitochondrial proteome is estimated to contain ∼1100-1500 nuclear-encoded proteins (Rath et al., 2021). To focus our efforts, we selected genes conserved in flies, humans, and mice that have been reported to localize to mitochondria (Lotz et al., 2014; Öztürk-Çolak et al., 2024; Rath et al., 2021). Of these, 256 genes had publicly available fly lines from the Transgenic RNAi Project (TRiP) collection and could be expressed in germline tissues (Perkins et al., 2015) (Figure 1A, Supplemental Table 1). Whenever possible, we independently tested multiple RNAi lines targeting the same gene.

**Fig 1.**
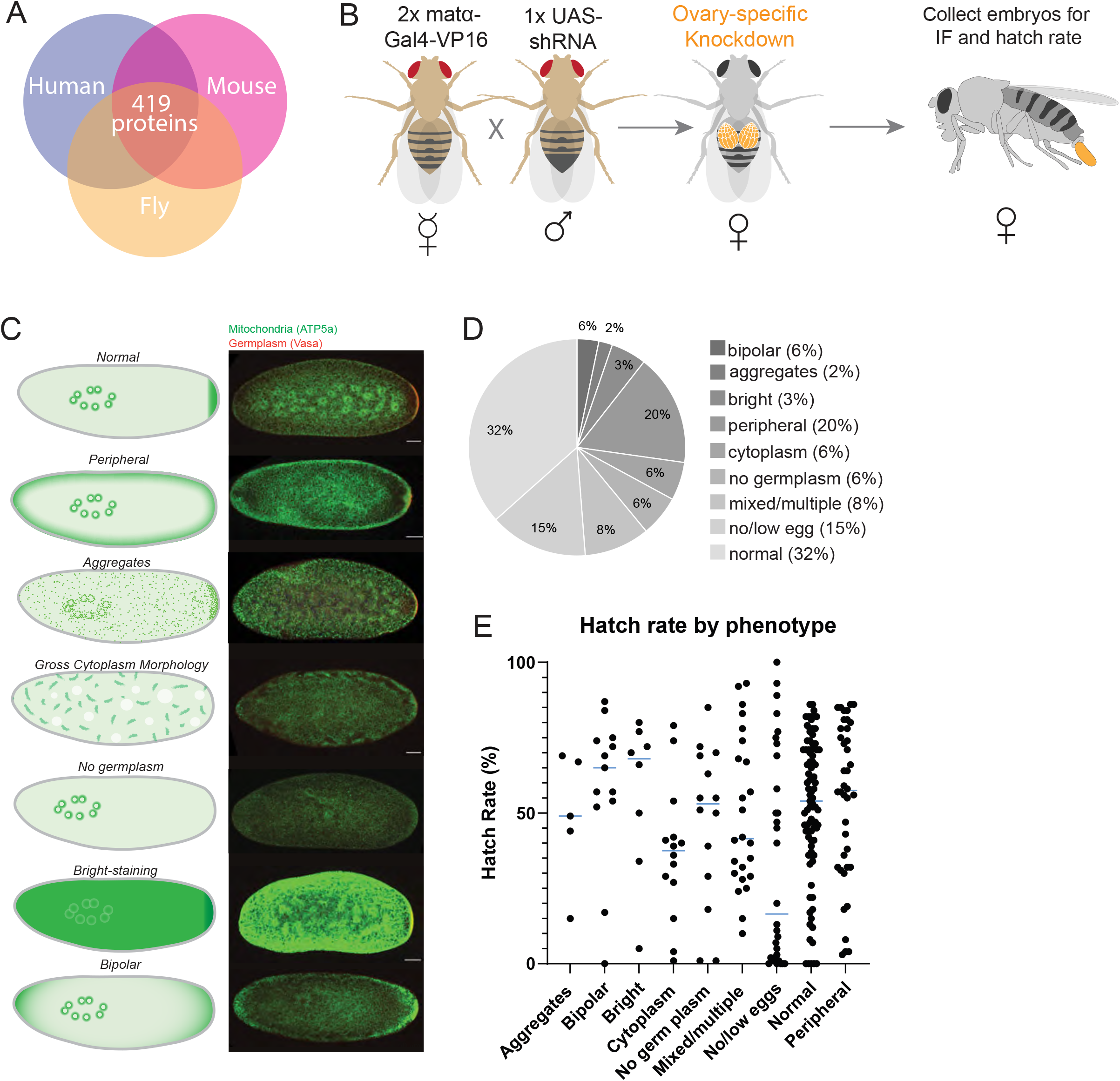
Maternal mitochondrial protein knockdown results in distinct embryo phenotypes. A. Venn diagram representing proteins associated with mitochondria in humans, mice, and flies. At least 419 proteins are conserved and exhibit mitochondrial localization across all three species. B. Cartoon schematic of crossing scheme. Female flies homozygous for two separate insertions of the oogenesis-specific driver, *matα-Gal4*, were crossed to male flies carrying the *UAS*-*shRNA*. Females F1 offspring with two copies of the *matα -Gal4* driver and one copy of *UAS*-*shRNA* were selected for embryo collections. Embryos were fixed and immunostained or incubated for an additional 30 hours to determine hatch rates. C. Left: cartoon representations of phenotypic classes observed by immunostaining. Relative intensity of green color indicates relative concentration of mitochondria. Right: example 20x confocal images of a syncytial stage embryo from each phenotypic class. Anti-ATP5α, marking mitochondria, is shown in green; anti-Vasa, marking the germplasm, is shown in red. Embryos are oriented with posterior end on the right. Scale bar: 50 µm. D. Pie chart showing relative frequency of each phenotypic class. Segments correspond to the classes shown in (C), with the addition of the ‘no/low egg’ class and the ‘mixed/multiple’ class, which showed multiple phenotypes but none claiming a majority. E. Hatch rate analysis by phenotypic class. Each dot represents the cumulative hatch rate for a single TRiP line. Blue line indicates mean value for each class. Associated data is shown in Table 1.

To achieve germline-specific gene disruption, we used the UAS-Gal4 system to bring together *UAS*-controlled shRNAs with an oogenesis-specific driver, *matα-Gal4* (Brand & Perrimon, 1993). Crossing these flies produced F1 offspring with two copies of the *matα-Gal4* driver and one copy of *UAS*-shRNA. This ensured specific expression of the shRNA constructs during mid-to-late oogenesis (stages 5-14) in F1 females and a strong reduction of the target RNAs in their offspring (Figure 1B). 0-2 hr old embryos were collected from F1 females, and mitochondria were visualized by immunofluorescence using an antibody against ATP5α, a component of Complex V of the electron transport chain that localizes to the inner mitochondrial membrane. We probed for DNA to assess mitotic progression and germplasm to mark the posterior end. Using laser scanning confocal microscopy, we imaged embryos at 20x magnification and examined the distribution and morphology of mitochondria (Supplemental Figure 1). In wild-type embryos, mitochondria either associate with dividing syncytial nuclei, concentrate in the posterior germplasm, or are homogeneously distributed throughout the embryo cytoplasm. We categorized each RNAi line into a single phenotypic category based predominantly on mitochondrial staining and identified distinct phenotypes (Figure 1C, D).

Among the 256 TRiP lines tested, we identified nine phenotypic classes: 32% (89 lines targeting 85 genes) showed no obvious disruption to the mitochondrial network and were considered “Normal”. Embryos with disruptions in the spatial distribution of mitochondria accounted for an additional 32% of lines (targeting 66 genes) and were divided into three categories: “No germplasm”, marked by a loss of posterior enrichment that could be explained by an upstream loss of germplasm assembly, “Bipolar”, showing both anterior and posterior mitochondrial enrichment, and “Peripheral”, showing mitochondria concentrating along the entire perimeter of the embryo cortex. Approximately 6% (14 lines targeting 11 genes) produced embryos with gross morphological disruptions in mitochondrial organization within the cytoplasm and defects in nuclear cycling/migration, termed “Cytoplasm”. Knockdown of five genes (2% of lines) produced embryos with “Aggregates”, suggesting defects in mitochondrial network dynamics. An unexpected “Bright” phenotype was observed in nine lines (3%) targeting seven genes, marked by extremely high ATP5α staining intensity. Approximately 8% of lines produced embryos with inconsistent or partially penetrant phenotypes, termed “Mixed/multiple” as they exhibited multiple phenotypes with none claiming a majority. Notably, ∼15% of lines were classified as “No/low eggs” either due to egg-laying defects, indicating a failure to complete oogenesis, or to low sample yield during fixation and devitilination, which can result from structural defects in the egg; thus, their effect on mitochondrial function in the embryo could not be examined. We included several positive controls to ascertain knockdown efficiency and to assist in defining the phenotypic classes: for example, Tropomyosin II (required for germplasm assembly) (Erdélyi et al., 1995) as a control for “no germplasm” and Mitofusin (a mediator of outer mitochondrial membrane tethering and fusion) (Bliek et al., 2013) for “aggregates”. Both lines were penetrant at nearly 100%, indicating a high knockdown efficiency.

We analyzed hatch rates for each RNAi line to determine whether viability correlated with any particular phenotype. We observed a wide range of hatch rate frequencies, even within the class that appeared phenotypically normal by mitochondrial staining (Figure 1E, Supplemental Table 1). Neither mitochondrial phenotypic class nor defect severity predicted hatch rate: some embryos with disorganized mitochondria hatched, whereas some with phenotypically normal mitochondria failed to complete embryogenesis. Morphological defects and embryonic viability are thus largely orthogonal readouts of mitochondrial function, supporting the need to consider both cellular and functional (i.e., metabolic) consequences to assess the maternal contribution to mitochondrial inheritance.

### Sideroflexin 1/3 influences mitochondrial size in embryos

Because of its striking and unexpected phenotype, we selected the ‘Bright’ phenotypic class to explore its origin in mitochondrial biology. Nine TRiP lines targeting seven genes led to the Bright phenotype (Supplemental Table 1, Supplemental Figure 2). These embryos appeared ∼10-fold brighter by ATP5α antibody staining than embryos in any other phenotypic category. Among the genes in this category, we focused on Sideroflexin 1/3, a mitochondrial inner membrane protein proposed to transport amino acids (Block et al., 2026; Fitzpatrick et al., 2026; Kory et al., 2018). *D. melanogaster* Sideroflexin 1/3 (*sfxn1-3*) is the sole homolog of the mammalian Sideroflexin 1 and 3 genes, and is hereafter referred to as ‘Sfxn’. Maternal Sfxn depletion produced a consistently strong and highly penetrant embryonic phenotype (nearly 100% of embryos collected). We ruled out trivial explanations for the observed phenotype, including reagent artifacts, antigen overexpression, and mitochondrial overproliferation (Supplemental Figure 3). Importantly, the bright staining was specific to epitopes inside the mitochondrial matrix, as it was also observed with an antibody to a different subunit of complex V, ATP5β, but not to an HA affinity tag targeted to the outer mitochondrial membrane (‘MitoTag’, see below) or to Vasa, which is cytoplasmic (Supplemental Figure 3). Together, these observations suggest that the bright phenotype may be reporting on a mitochondrial alteration that permits increased antigen detection or accessibility.

We reasoned that Sfxn knockdown may affect mitochondrial morphology and therefore examined embryos using high-resolution Airy scan imaging (Figure 2A, B). In control embryos (expressing shRNA to a nontargeting control sequence), the mitochondrial network was highly fragmented and homogeneous; individual mitochondria appeared spherical, with an approximate diameter of <0.5 μm (Figure 2A). By contrast, in embryos where *sfxn1-3* RNA was targeted maternally (hereafter referred to ‘Sfxn RNAi’ embryos) mitochondria appeared strikingly enlarged and highly variable with some mitochondria approaching several microns in diameter (Figure 2B). However, because ATP5α, which normally concentrates at cristae tips, labeled these structures inconsistently, and light microscopy cannot resolve closely apposed organelles, we could not determine whether the ATP5α-positive structures were single enlarged mitochondria or tight clusters of smaller ones.

**Fig 2.**
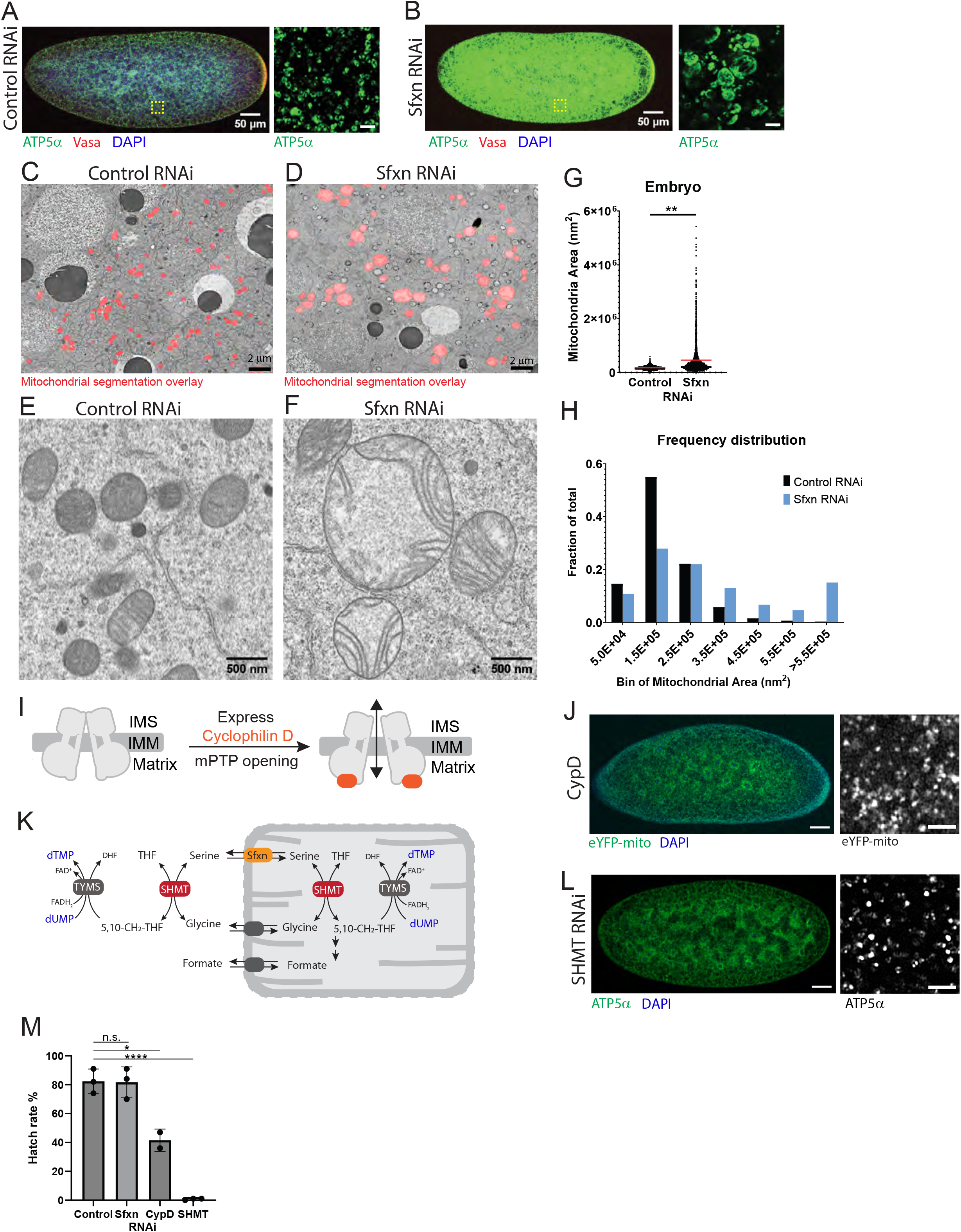
Sideroflexin 1/3 RNAi results in enlarged but intact mitochondria. A-B. Immunofluorescence images of syncytial stage (A) control RNAi or (B) Sfxn RNAi embryos. Left: 20x laser scanning confocal image. Scale bar: 50 μm. Right: 63x airyscan image of mitochondria. Scale bar: 2 μm. C-F. Electron microscopy (EM) images of syncytial-stage embryos. Example images are from (C and E) control RNAi or (D and F) Sfxn RNAi embryos. (C-D) EM images overlayed with results from mitochondrial segmentation shown in red. Scale bar: 2 μm. (E-F) Scale bar: 500 nm. G-H. Quantification of mitochondrial area from STEM images shown as (G) violin plot or (H) histogram. Results are from n = 3 embryos from each condition. Control: n = 6686 mitochondria. Sfxn RNAi: n = 2677 mitochondria. **p value = 0.006, two-tailed nested t-test. I. Cartoon schematic showing proposed mechanism of mitochondrial permeability transition pore (mPTP) opening upon cyclophilin D (CypD) binding. J. Left: 20x image of an embryo expressing human cyclophilin D. Mitochondria are shown in green, marked by endogenously expressed eYFP-mito. DNA, marked by DAPI, is shown in blue. Scale bar: 50 μm. Right: Single channel 63x image of mitochondria from the embryo shown on the left. Scale bar: 2 μm. K. Schematic depicting serine hydroxymethyltransferase (SHMT)-dependent steps in one-carbon metabolism and thymidylate synthase (TYMS)-dependent dUMP to dTMP conversion. L. Example immunofluorescence images of a SHMT-depleted early embryo. Left: 20x confocal image. Green: ATP5α; red: Vasa; DNA (DAPI). Scale bar: 50 µm. Right: Single channel 63x image of mitochondria from the embryo shown on the left. Scale bar: 2 μm. M. Bar chart of hatch rate analyses. Each dot represents an independent hatch rate experiment. Hatch rate for control RNAi (82.3%, n = 906 embryos) and Sfxn RNAi (81.7%, n = 1159 embryos) was not significantly different (p = 0.94). Hatch rate for CypD-expressing embryos (41.5%, n = 959 embryos) was significantly lower than control (p = 0.02), as was the hatch rate for *Shmt* RNAi embryos (0.7%, n = 1067 embryos, ****p <0.0001). p values calculated by two-tailed unpaired t-test.

To examine mitochondrial morphology at the nanometer scale, we turned to electron microscopy (EM). In control embryos, mitochondria appeared rounded, electron-dense, and exhibited ordered cristae in nearly all detected mitochondria (Figure 2C, E). To quantify mitochondrial size in control and Sfxn-depleted samples, we applied the deep-learning-based model MitoNet to segment mitochondria in our EM datasets (Conrad & Narayan, 2023). In control embryos, organelle-to-organelle variability in size and shape was low, with an average mitochondrial cross-sectional area of 0.177 +/− 0.087 *µ*m^2^ (n = 6686 mitochondria, +/− S.D) (Figure 2G, H). Compared to control, mitochondria in Sfxn RNAi embryos were ∼2.3-fold larger, on average, and were also much more variable in size (0.418 +/− 0.582 *µ*m^2^, n = 2677 mitochondria, +/− S.D) (Figure 2D, F-H). Notably, these enlarged mitochondria typically lacked evidence of damage or degradation: they had intact outer and inner membranes and maintained lamellar cristae organization (Figure 2F). In addition, whereas the largest mitochondria in the control samples appeared to be undergoing fission, the largest mitochondria in the Sfxn-depleted samples were always spherical (Supplemental Figure 4). Excessive mitochondrial matrix swelling stretches the outer mitochondrial membrane and can lead to bursting (Sesso et al., 2012); however, we observed no evidence of mitochondrial membrane herniation (n = 2677 mitochondria). Together, these results show that mitochondria in Sfxn-depleted embryos are isotropically enlarged but maintain their structural integrity.

### Sideroflexin affects mitochondrial morphology independent of known regulators of mitochondrial membrane permeability and dynamics

We next explored whether the mitochondrial swelling observed in Sfxn RNAi embryos resulted from known mechanisms that disrupt mitochondrial integrity, such as alterations to mitochondrial inner membrane permeability or dynamics.

Opening of the mitochondrial permeability transition pore (mPTP), a nonspecific pore in the inner membrane, permits solute entry and causes osmotic swelling of the matrix (Bernardi et al., 2023). We therefore asked if perturbations that stimulate mPTP formation would phenocopy the morphology we observe upon Sfxn depletion. To force activation of the mPTP, we expressed human cyclophilin D (CypD), a regulator of mPTP opening (Figure 2I). CypD expression in the *D. melanogaster* ovarian germline stem cells (e.g. the earliest stages of oogenesis) produces germ cells with swollen mitochondria containing fragmented cristae, ultrastructural features consistent with mPTP opening (Monteiro et al., 2023). We expressed human CypD (*UASp*-CypD-HA) maternally under *matα-Gal4* control and collected 0–2 hr old embryos. Using eYFP-mito to visualize mitochondria, which reports on mitochondrial morphology similarly to the ATP5α immunostaining (Supplemental Figure 5), we found that CypD-expressing embryos were indistinguishable from control with no evidence of enlargement (Figure 2J). However, CypD-expressing embryos did show a significantly reduced hatch rate relative to control (41.5% for CypD versus 82.3% for control, p = 0.02), indicating that forced CypD expression produced a pathological and partially lethal phenotype in the germline (Figure 2M). By contrast, Sfxn depletion had little effect on embryo survival (81.7%, p = 0.94 compared to control). CypD expression therefore compromises embryogenesis without the morphological hallmarks of Sfxn loss. Thus, the presence of intact inner and outer membranes, preserved lamellar cristae, and a complete absence of mitochondrial herniation in Sfxn RNAi embryos suggest that mitochondrial enlargement caused by Sfxn loss is independent of the mitochondrial permeability transition pore.

Next, we compared the Sfxn RNAi phenotype with that of genes regulating mitochondrial network dynamics. Loss of the mitochondrial fission genes, such as Drp1 and Marf/Mitofusin increases mitochondrial size by increasing connectivity, leading to elongated strands of fused mitochondria (Chowdhary et al., 2020). The Sfxn knockdown phenotype is strikingly different as it causes isotropic enlargement of the organelle. Indeed, fission/fusion regulators, including Marf/mitofusin were identified as members of the “aggregate” phenotypic category in our RNAi screen (Figure 1C and Supplemental Table 1), resembling the mitochondrial defects described in other tissues (e.g., neurons) (Rikhy et al., 2007). These lines also lacked the ‘bright’ phenotype and caused near-complete embryonic lethality (Chowdhary et al., 2020). Together, our data show that loss of Sfxn results in a mitochondrial morphological phenotype distinct from that resulting from disrupted fission/fusion dynamics.

#### Mitochondrial swelling in Sfxn-depleted embryos is not due to a deficiency in one-carbon metabolism

Mammalian Sideroflexin 1 was initially identified as a serine importer in a screen sensitized to dependency on the mitochondrial one-carbon pathway (Kory et al., 2018). Serine delivered by this route is used by mitochondrial serine hydroxymethyltransferase (SHMT) to catalyze the reversible conversion of serine into glycine, generating activated one-carbon units (5,10-methylene-THF) needed for cell proliferation (Figure 2K). If serine delivery to mitochondria was also the principal function of Sfxn in the *Drosophila* germline, Sfxn depletion and mitochondrial one-carbon deficiency should produce similar phenotypes.

*Drosophila* has only one *Shmt* gene, which provides both cytoplasmic and mitochondrial SHMT (Winkler et al., 2017). We expressed shRNA against *Shmt* during oogenesis and collected 0-2 hr old embryos laid by these mothers (hereafter referred to as ‘SHMT RNAi embryos’). We found that SHMT RNAi embryos immunostained with ATP5α neither exhibited the bright staining nor the mitochondrial swelling characteristic of Sfxn RNAi embryos (Figure 2L). Rather, SHMT RNAi embryos arrested at approximately nuclear cycle 13 and were completely inviable (hatch rate 0.7%, n = 1067 embryos) (Figure 2M). (Supplemental Figure 6) (Winkler et al., 2017). These data show that maternally supplied SHMT is not required for mitochondrial volume control. Depleting mitochondrial SHMT from mammalian cells also causes defects in mitochondrial respiration, in part because of its role in mitochondrial translation and electron transport chain (ETC) assembly (Minton et al., 2018; Morscher et al., 2018). We therefore tested ETC assembly and, specifically, Complex I function in Sfxn RNAi embryos. We found no change in electron transport chain assembly or Complex I function, suggesting that respiratory capacity is maintained upon maternal depletion of Sfxn (Supplemental Figure 6). Together, these observations indicate that Sfxn’s germline role is not explained by its contribution to mitochondrial one-carbon metabolism.

Given its prominent enlarged mitochondrial phenotype, we explored alternative models for Sfxn function. Consistent with its described role as a serine transporter, we hypothesized that Sfxn may regulate metabolite flux across the inner mitochondrial membrane. Furthermore, because mitochondrial swelling had not been observed in cultured mammalian cells (Kory et al., 2018), we speculated that Sfxn-dependent mitochondrial swelling could be developmentally regulated.

### Mitochondrial swelling occurs only during egg activation after Sfxn depletion

After Sfxn knockdown, mitochondria were enlarged in 100% of early embryos, even those as early as the first nuclear division (∼10-30 minutes after egg laying). To determine when during development mitochondrial morphology changes first appear in response to Sfxn RNAi, we examined egg chambers at different stages of oogenesis (Figure 3A). Because immunostaining oocytes can be challenging due to the eggshell and vitelline membrane, which impede penetration at late stages of oogenesis, we used a fly line that ubiquitously expresses a fluorescently tagged mitochondrial matrix marker, mito-eYFP (Lajeunesse et al., 2004). After confirming that mito-eYFP labels the same structures as ATP5α immunostaining (Supplemental Figure 5), we examined stage 14 oocytes, the final state of oogenesis. In mature oocytes expressing shRNA to a nontargeting control sequence (hereafter referred to as ‘control’ oocytes), mitochondria appear homogeneous and punctate, similar to how they appear in embryos (Figure 3B). Surprisingly, in mature oocytes expressing *Sfxn* shRNA (hereafter referred to as ‘Sfxn RNAi’ oocytes), mitochondria were largely indistinguishable from those in control oocytes (Figure 3C). The high density of mitochondria in oocytes made accurate quantification difficult; we therefore turned to transmission electron microscopy (Figure 3D-G). In control oocytes, mitochondria appeared electron-dense and circular in cross-section; occasionally, we observed figure-eight-shaped mitochondria, consistent with mitochondrial fission. Mitochondria were more clustered than in embryos and often separated by only a few nanometers (Figure 3D, E). The average area of a mitochondrion in control oocytes was 0.156 +/− 0.074 *µ*m^2^ (n = 2898 mitochondria, +/− S. D.), slightly smaller than in embryos (0.177 +/− 0.087 *µ*m^2^) (Figure 3H). Mitochondria in Sfxn RNAi oocytes were morphologically similarly shaped to those in control samples: circular in cross-section and occasionally figure-eight shaped; additionally, the mean area for these mitochondria was 0.199 +/− 0.081 µm^2^ (n = 1841 mitochondria, +/− S. D.), a statistically insignificant increase compared to the control (p = 0.102, two-tailed nested t-test) (Figure 3H). Importantly, mitochondria in Sfxn RNAi oocytes did not display the massive size variability observed in Sfxn RNAi embryos (Figure 3I). Because mitochondria did not appear enlarged at the latest stages of oogenesis but were enlarged at the earliest stages of embryogenesis (Figure 3J), we infer that mitochondrial swelling occurs during the egg-to-embryo transition.

**Fig 3.**
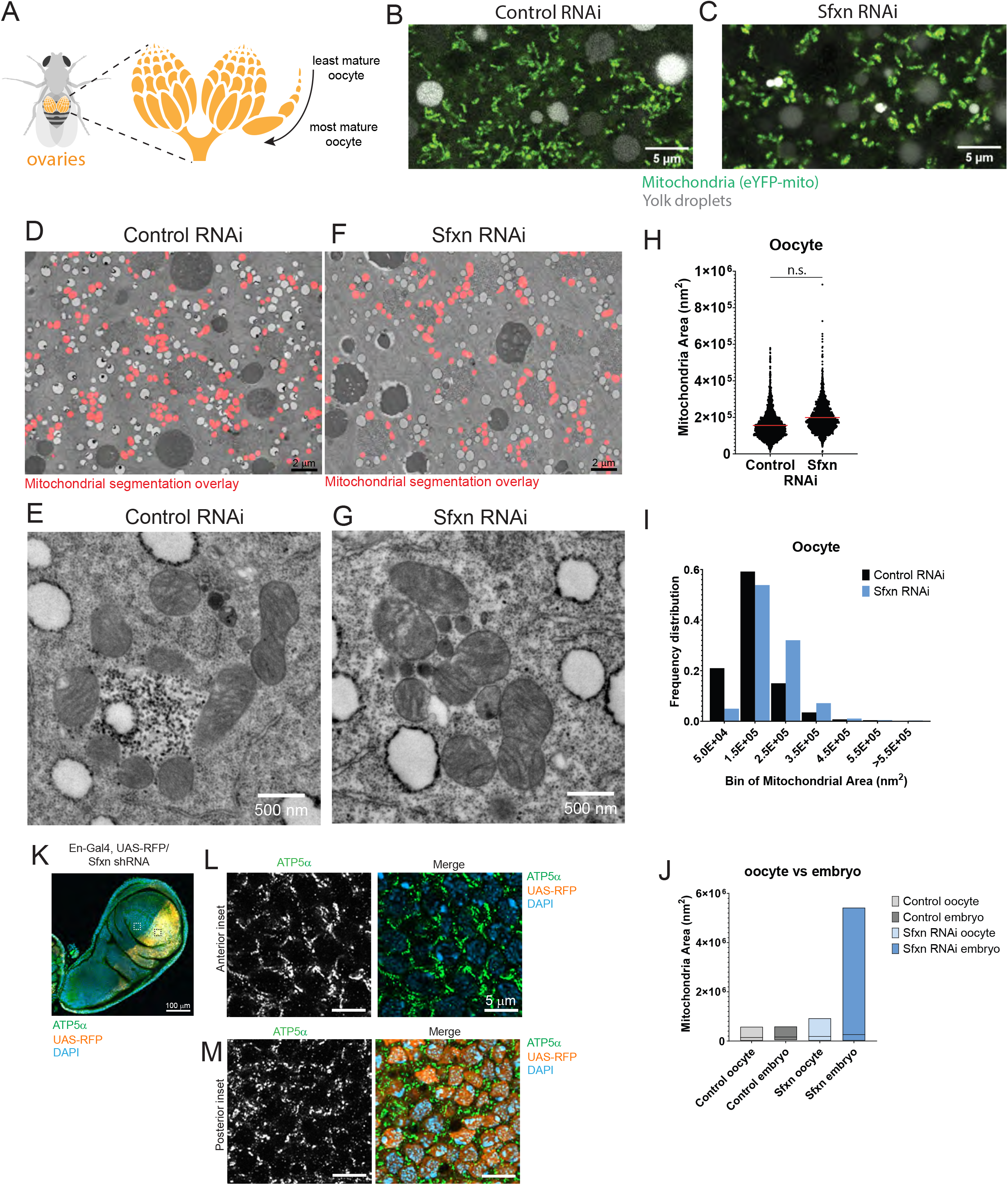
Mitochondrial swelling occurs during egg activation. A. Cartoon of female fruit fly with ovaries highlighted; magnification of ovary cartoon showing developmental progression of oocyte development within a single ovariole. B-C. Confocal image of mature (stage 14) oocyte from either (B) control or (C) Sfxn RNAi embryos expressing an eYFP-mitochondrial matrix marker. D-G. Scanning transmission electron microscopy (STEM) images of mature (stage 14) oocytes expressing either (D and E) control shRNA or (F and G) Sfxn RNAi. (D, F) STEM images overlayed with results from mitochondrial segmentation shown in red. H-I. Quantification of mitochondrial area from STEM images of oocytes in (H) violin plot or (I) histogram. Results are from n = 3 oocytes from each condition. Control RNAi: n = 2898 mitochondria. Sfxn RNAi: n = 1841 mitochondria. J. Comparison of mitochondrial area from oocytes and embryos. K-M. Laser scanning confocal images of late larval wing disc. *UAS*-driven red fluorescent protein (RFP) is expressed in the posterior compartment and marks the cells that express *UAS*-driven shRNA to *sfxn1-3*. Mitochondria marked by ATP5α is shown in green. DNA marked by DAPI is shown in light blue. K. 20x image showing whole wing disc with dotted-line boxes indicating insets shown in (L) and (M). L. 63x image of anterior compartment of wing disc (white dotted line inset from K). ATP5α channel is shown in greyscale next to three-color overlay image. M. Posterior compartment of wing disc (black dotted line inset from K). ATP5α channel is shown in greyscale next to three-color overlay image.

To test whether Sfxn’s role in mitochondrial morphology is unique to the egg-to-embryo transition, we depleted Sfxn in somatic tissues (Figure 3K-M). We chose the larval wing disc, where an *engrailed-Gal4* driver is expressed only in the posterior compartment, leaving the anterior compartment as an internal control to compare mitochondrial morphology. We identified the posterior cells by expression of a *UAS*-driven mCherry marker. When comparing the two compartments, we found that mitochondria appeared indistinguishable both in mitochondrial morphology and in ATP5α reactivity (Figure 3L, M). These results suggest that Sfxn-sensitive mitochondrial enlargement and ATP5α hyper-reactivity are specific to the egg-to-embryo transition.

### Sfxn prevents neutral amino acids from accumulating in mitochondria

The developmental timeline of mitochondrial swelling after Sfxn knockdown suggests a specific requirement for Sfxn during the egg-to-embryo transition, but it does not explain how loss of Sfxn function leads to this phenotype. It has been proposed that Sfxn family members, tested primarily in mammalian tissue culture, transport neutral amino acids into mitochondria, including serine (Kory et al., 2018), proline, glycine, taurine, hypotaurine, and GABA (Block et al., 2026) and leucine (Fitzpatrick et al., 2026). Given the breadth of potential amino acid substrates, we directly asked whether Sfxn depletion affected amino acid abundance or distribution at the egg-to-embryo transition. To do so, we performed targeted metabolomics analysis using hydrophilic interaction liquid chromatography-mass spectrometry (HILIC-MS). We focused on polar metabolites, which would allow robust detection of putative Sfxn substrates.

To determine the steady-state metabolic profile of tissues before and after the egg-to-embryo transition, we analyzed whole-tissue lysates from isolated stage 14 oocytes and early embryos (0-2.5 hr old). Of the 221 metabolites in our target list (Supplemental Table 2), ∼200 metabolites were consistently detected across all whole-tissue lysate experiments. Analysis of whole stage 14 oocyte lysates revealed a largely unchanged metabolome in Sfxn RNAi oocytes compared to control (Figure 4A, B). The majority of detected metabolites, including free proteinogenic amino acids, showed no or modest changes (FC range 0.73–1.95) (Figure 4B). These results indicate that the total amino acid pool was preserved in Sfxn RNAi oocytes at the whole-cell level. Similarly, in whole embryo lysates, we again observed a largely unchanged metabolome when comparing Sfxn-depleted samples to control (Figure 4C, D). Metabolite levels were similar between Sfxn RNAi and control, and between whole oocyte lysate and embryos, except for one notable difference: we observed a pronounced and reproducible accumulation of beta-alanine across three biological replicate experiments (FC = 10.23 +/− 0.82, mean +/− S.D.; Figure 4C, D). Beta-alanine is a polar, uncharged amino acid required for carnosine synthesis and a putative substrate of human Sfxn (Block et al., 2026). In contrast, total serine levels were not significantly altered in Sfxn RNAi embryos, despite robust detection (FC = 1.39 +/− 0.13, mean +/− S.D.). Glycine, the coproduct of mitochondrial SHMT-mediated serine catabolism, was well-detected but only marginally elevated in Sfxn RNAi (FC = 1.51, +/− 0.27). As a control, we also performed metabolomics on embryos maternally depleted of SHMT. Consistent with our finding that SHMT RNAi embryos did not phenocopy the mitochondrial swelling conserved in Sfxn RNAi embryos (Figure 2L), metabolomics on these embryos similarly revealed notable differences. Unlike in Sfxn RNAi embryos, SHMT RNAi embryos displayed a clear signature of global one-carbon deficiency consistent with impaired thymidylate synthase (TS) activity, which requires methylene-THF as a one-carbon donor to convert deoxyuridine monophosphate (dUMP) to deoxythymidine monophosphate (dTMP) (Figure 2K). Extracts from SHMT RNAi embryos showed a reduction in dTMP and an elevation in dUMP, free deoxyuridine (a catabolite of dUMP), and free uracil (Supplemental Figure 4). The absence of these hallmarks in Sfxn RNAi embryos indicates that global one-carbon metabolism was apparently preserved, either because the cytosolic SHMT branch can compensate when serine flux through mitochondria is impaired or because the germline’s reliance on the mitochondrial arm of the one-carbon pathway is already low. Together, these data suggest that Sfxn function does not contribute to total metabolite abundance in mature oocytes or early embryos.

**Fig 4.**
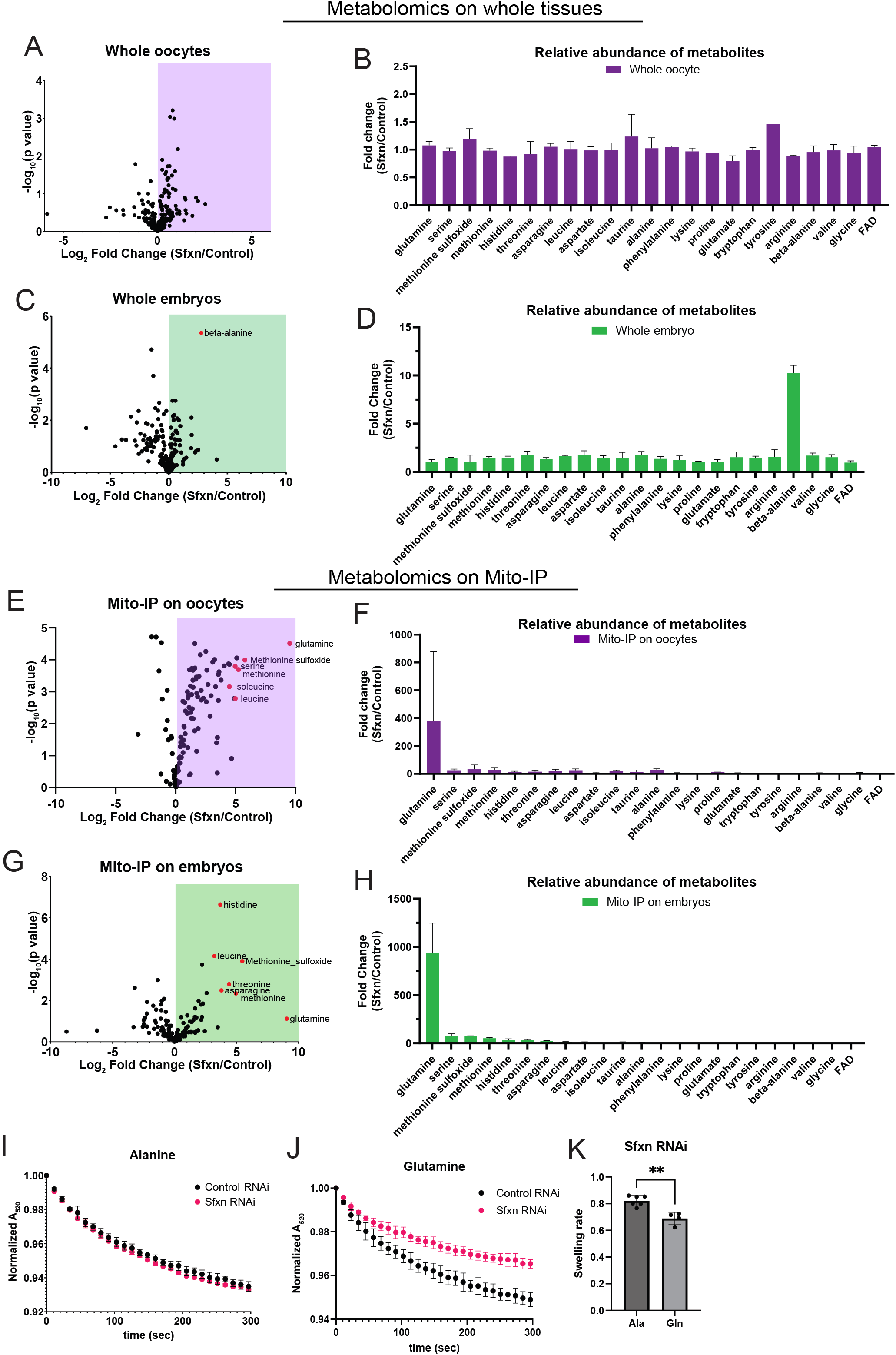
Neutral amino acids accumulate in Sfxn-deficient mitochondria from both oocytes and embryos. A-D. Metabolomics on whole tissue lysates from oocytes or embryos. E-H. Metabolomics on immunopurified mitochondria. A, C, E, D. Example volcano plots from the indicated experiment. B, D, F, H. Relative abundance of indicated amino acids and metabolites. All measurements are metabolite abundance in Sfxn RNAi samples relative to control RNAi samples. Mean +/− S.D. Embryo samples are from syncytial stage (aged 0-2 hr old). Oocyte samples are from mature (stage 14) oocytes. I-J. Swelling assay curves of isolated mitochondria from either control or Sfxn RNAi embryos treated with (I) alanine or (J) glutamine. Values are normalized to absorbance at t = 0. n = 3 technical replicates per curve, ± SD. K. Bar chart summary from swelling assays showing relative swelling rate of mitochondria isolated from Sfxn RNAi embryos relative to control for alanine (ala) and glutamine (gln)

Given the reported function of Sfxn family members in mitochondrial amino acid transport, we next asked whether Sfxn impacts amino acid abundance specifically in mitochondria, which may have been obscured in whole-cell lysate samples. We performed mitochondrial immunoprecipitation (mito-IP) on oocyte and embryo samples stage-matched to their whole-tissue counterparts. We adapted the ‘MitoTag’ approach, which utilizes a mitochondrially targeted 3xHA epitope tag for rapid mitochondrial isolation (W. W. Chen et al., 2017), to generate constructs for tissue-specific expression in flies using the UAS-Gal4 system (Supplemental Figure 7). We generated two independent mitochondria-targeting constructs: a TOM20-based HA-tagged anchor (MitoTag1) and an OMP25-based HA-tagged anchor (MitoTag2) under *UAS* control (Supplemental Figure 7). Samples were collected from females expressing the MitoTag construct in late oogenesis via the *Matα-Gal4* driver. We verified the effectiveness of our mitochondrial isolation strategy by confirming that metabolites known to be excluded from mitochondria, including cystine (enriched in lysosomes), glucose, and sedoheptulose-7-phosphate (cytoplasmic), were not present in mito-IP fractions above background while being robustly detected in whole tissue lysates (W. W. Chen et al., 2016). Because both lines showed comparable metabolite detection and abundance, we used the Omp25-based version for consistency with prior literature.

Strikingly, when we examined the metabolic profile of isolated mitochondria, we observed a massive accumulation of free amino acids both in Sfxn RNAi oocyte (Figure 4E, F) and embryo (Figure 4G, H) samples compared to control. The most enriched metabolite by an order of magnitude was glutamine, which accumulated to extremely high levels in Sfxn RNAi mitochondria in both stages (Figure 4E-H). Although somewhat variable, glutamine abundance inside the mitochondrial compartment increased nearly 1000-fold in Sfxn-depleted samples relative to control (383.0 +/− 495.0 in oocytes, 938.6 +/− 308.8 in embryos; mean FC +/− SD). The next most enriched amino acid was serine, a proposed Sfxn substrate (21.9 +/− 13.0 in oocytes, 77.2 +/− 20.3 in embryos; mean FC +/− SD). Methionine, histidine, threonine, asparagine, leucine, and isoleucine also showed enrichment across replicates, whereas lysine, tryptophan, tyrosine, and arginine showed little change (Supplemental Figure 8). FAD+, a mitochondrial-specific cofactor important for respiratory activity, was also not substantially changed. Several of these polar metabolites are consistent with the established or proposed substrate specificity of Sfxn (Block et al., 2026; Fitzpatrick et al., 2026; Kory et al., 2018), although we cannot rule out that mitochondrial accumulation of certain amino acids reflects indirect consequences of Sfxn loss.

The metabolite that exhibited the most substantial enrichment in Sfxn RNAi mitochondria, both in oocytes and in embryos, was glutamine. If this accumulation reflects a loss of Sfxn activity that reduces mitochondrial glutamine permeability, then mitochondria lacking Sfxn should show reduced glutamine flux across the inner membrane. To test this, we isolated mitochondria from freshly laid embryos and measured amino acid uptake kinetics using an *in vitro* swelling assay. This technique has been used to characterize the permeability of the mitochondrial inner membrane to a wide range of solutes (Beavis et al., 1985; Chappell, 1968; Garlid & Beavis, 1985; LaNoue & Schoolwerth, 1979). This light scattering assay takes advantage of the relationship between matrix volume and the amount of light scattered by a suspension of mitochondria. Compared to alanine, which was not substantially enriched in Sfxn RNAi embryo samples, elicited a similar swelling profile in control and Sfxn RNAi conditions (Figure 4I), glutamine influx was significantly reduced in mitochondria isolated from Sfxn-depleted embryos (Figure 4J, K). These data show that the reduction in glutamine permeability is solute-selective rather than a general loss of inner membrane integrity. Together, these data show that mitochondria in Sfxn RNAi tissues exhibit elevated amino acid levels, supporting a role for Sfxn in amino acid export.

### Sfxn is required for selective amino acid permeability of the mitochondrial inner membrane

While loss of maternal Sfxn function causes amino acids to accumulate equally in oocytes and embryos, mitochondrial volume changes only in the embryo. This decoupling of metabolic and morphological phenotypes suggests that a change in cellular physiology is required to trigger mitochondrial swelling. In flies, the egg-to-embryo transition involves a significant physiological transition through consecutive but distinct processes: ovulation, egg activation, and fertilization (Avilés-Pagán & Orr-Weaver, 2018). Ovulation triggers egg activation, which hinges on calcium signaling cascades that link cell cycle control, translational activation, and metabolic rewiring. Upon fertilization, the paternal and maternal nuclei fuse and rapid embryonic nuclear divisions begin (Figure 5A). To determine which process is sensitive to polar amino acid levels in mitochondria, we independently manipulated each step and analyzed mitochondrial morphology upon Sfxn knockdown.

**Fig 5.**
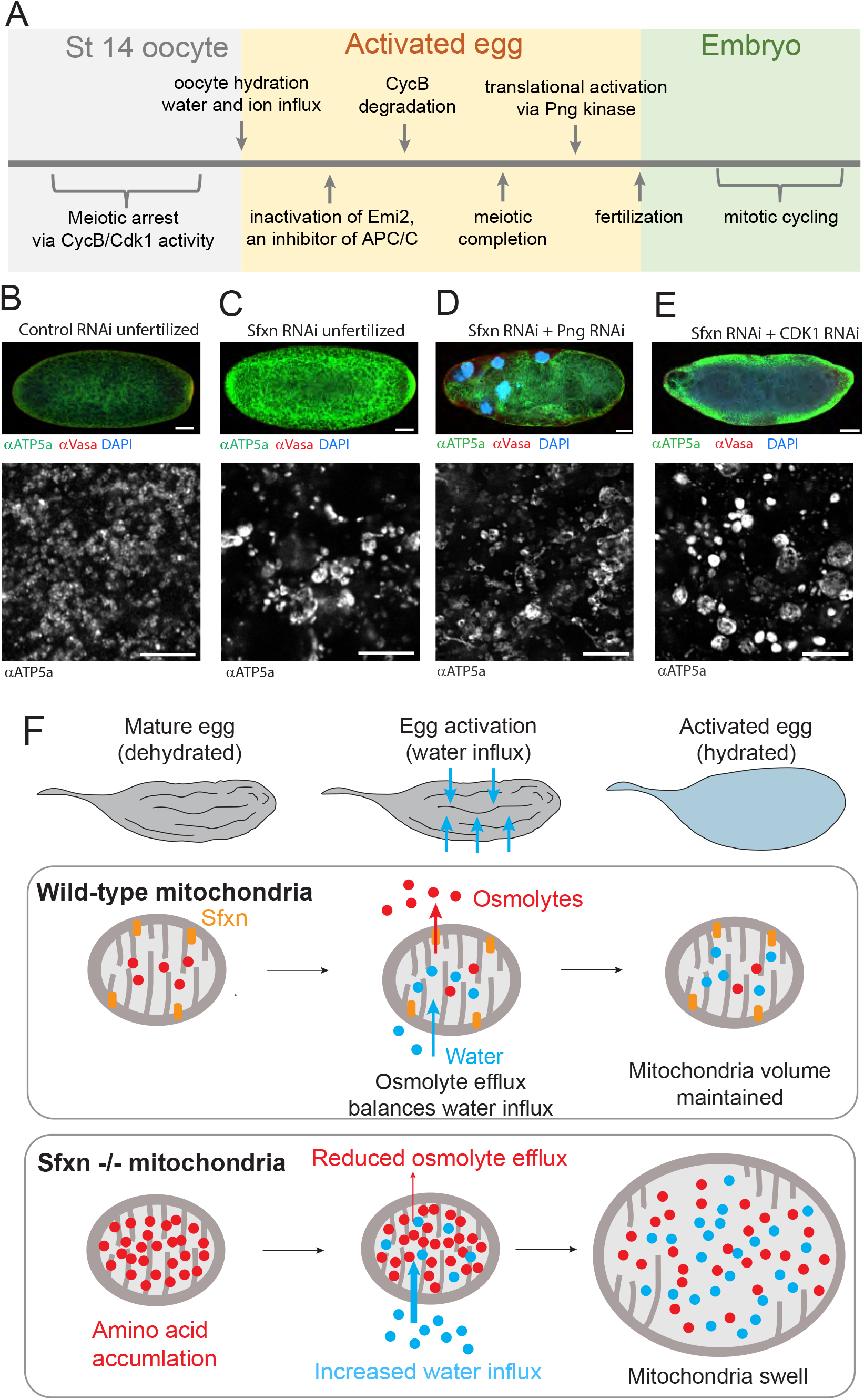
Mitochondrial volume homeostasis during oocyte hydration requires Sfxn. A. Timeline of key steps during egg activation. B-E. Test for epistasis of developmental events (outlined in A) with respect to Sfxn function. Top: Three-channel overlay images of control and experimental conditions. Green: ATP5α; Red: vasa; Blue: DAPI staining of DNA. Scale bar: 50 μm. Bottom: Single channel 63x images of mitochondria in the embryos shown above. ATP5α staining is shown in greyscale. Scale bar: 5 μm. F. Model for Sfxn function in the germline. During egg activation, dehydrated mature oocytes uptake extracellular water, causing the cytoplasm to swell. Mitochondria, in equilibrium with the cytoplasm, release neutral amino acids to accommodate the uptake of water during egg hydration. This equilibration depends on Sfxn function. When Sfxn is absent, osmotically active amino acids accumulate in the mitochondrial matrix, leading to swelling during egg hydration.

To determine whether fertilization-dependent mitotic cycling is required for mitochondrial swelling, we crossed F1 females (expressing shRNA against Sfxn in oocytes) with sterile males, thereby bypassing the mating requirement for normal egg-laying behavior. Mitochondria in control unfertilized eggs were similar to those in early embryos (Figure 5B). However, in unfertilized eggs with Sfxn depleted, mitochondria immunostained with the ATP5α antibody appeared bright and enlarged, similar to what we observed in fertilized, Sfxn-depleted embryos (Figure 5C). This shows that fertilization is not required for the Sfxn phenotype to manifest, indicating that mitochondrial swelling must occur prior to fertilization.

To target the earlier egg activation program directly, we generated double knockdowns of Sfxn together with Pan Gu kinase (Png) or Cdk1. Png triggers translational activation of maternal transcripts at egg activation, including Cyclin B, while Cdk1 drives cell cycle re-entry in meiotically arrested oocytes (Avilés-Pagán & Orr-Weaver, 2018; Berg et al., 2024). In both Sfxn; Png and Sfxn; Cdk1 double knockdown embryos, mitochondria remained bright and enlarged similar to Sfxn knockdown alone (Figure 5D, E). We confirmed that Png was sufficiently reduced in this background by the presence of giant nuclei, which form when limited Png activity causes endoreduplication of embryonic nuclei without nuclear division (Figure 5D, top). Similarly, we confirmed Cdk1 disruption by the absence of syncytial nuclei (besides the single pronucleus, a product of meiosis) (Figure 6C, top). Thus, Sfxn-dependent sensitivity to mitochondrial swelling precedes Png-dependent translational activation and Cdk1-dependent cell cycle re-entry.

Taken together, mitochondrial swelling in Sfxn-depleted eggs requires neither fertilization, nor Png-dependent translational activation, nor Cdk1-dependent cell cycle re-entry. By exclusion, we postulate that the trigger therefore occurs in parallel to egg activation or to ovulation, the trigger preceding it. During ovulation, rapid hydration of the oocyte is accompanied by an abrupt fall in cytoplasmic osmolarity (Horner & Wolfner, 2008a; Mahowald et al., 1983; York-Andersen et al., 2021). Mitochondria carrying an elevated matrix solute load would face a steeper inward osmotic gradient under these conditions and, lacking Sfxn, would have no route to release the accumulated amino acids (Figure 5F). We propose that mitochondrial swelling observed in Sfxn-depleted embryos results from the combination of a pre-existing solute burden and the osmotic shift associated with egg hydration.

## Discussion

Our study provides evidence that Sfxn functions in the germline to preserve mitochondrial morphology during the osmotic stress of egg hydration. Because mitochondrial inner membrane permeability largely determines matrix solute content, and because free amino acids are used by cells as biological osmolytes, amino acid content within the matrix is simultaneously a metabolic and biophysical parameter. We find that by controlling the concentration of free amino acids in mitochondria during oogenesis, Sfxn controls mitochondrial susceptibility to volume changes resulting from egg hydration during ovulation.

Why are oocyte and embryo mitochondria morphologically distinct after Sfxn knockdown if their metabolite composition is not? We propose that these morphological differences reflect differences in osmolarity across the inner mitochondrial membrane. Accumulation of amino acids would elevate osmotic pressure in the mitochondrial matrix and create a stronger driving force for water influx upon egg hydration, when the surrounding cytoplasmic osmolarity drops (estimated to be ∼200 mOsm in wild-type eggs) (Horner & Wolfner, 2008a). The abruptness of this osmotic shift precipitates the overt mitochondrial swelling observed in Sfxn RNAi embryos and may explain why swelling is absent in dehydrated, mature oocytes.

Three observations support an osmotic trigger rather than a signaling one. First, mitochondrial swelling requires neither fertilization and the mitotic divisions that follow it, nor Png-dependent translational activation, nor Cdk1-dependent cell cycle re-entry; disrupting either does not prohibit mitochondrial swelling, placing the trigger upstream of, or in parallel with, the known signaling outputs of egg activation. Second, the phenotype is confined to the germline, since Sfxn depletion in the larval wing disc produces neither enlargement nor elevated ATP5α reactivity, consistent with a requirement that arises only in a cell facing this particularly abrupt osmotic transition. Third, the enlarged mitochondria retain intact inner and outer membranes and lamellar cristae and show no herniation, and expression of human cyclophilin D does not reproduce the enlargement, arguing that the swelling is osmotic rather than a consequence of permeability transition.

Our interpretation of Sfxn function relies on established properties of the organelle. Solute permeability across the mitochondrial inner membrane is the primary barrier to osmotic regulation because the outer membrane is freely permeable to small solutes, whereas the inner membrane is not (Cunningham & Rutter, 2020; Varughese et al., 2021). Furthermore, isolated mitochondria behave as near-ideal osmometers, capable of reversibly adjusting their matrix volume in response to changes in the solute composition of the external media (Bentzel & Solomon, 1967; Tedeschi & Harris, 1955). Matrix volume is itself a regulated parameter held within a narrow range by K⁺/H⁺ cycling and other volume-regulatory transport (Garlid & Paucek, 2003; Halestrap, 1989). Finally, the solutes cells use to buffer osmotic stress in the cytoplasm are largely free amino acids and their derivatives: taurine, glycine, proline and β-alanine are canonical compatible osmolytes whose efflux is a core effector of regulatory cell volume decrease (Hoffmann et al., 2009; Yancey, 2005; Yancey et al., 1982). These amino acids are among those that accumulate in Sfxn-depleted mitochondria.

Prior studies of Sfxn function were designed to detect disruptions in metabolite permeability using sensitized screens or uptake assays, which reveal import capacity but cannot directly report on export or physiological transport functions (Block et al., 2026; Fitzpatrick et al., 2026). A system designed to uncover an export defect would require conditions in which the cytoplasm depends on mitochondrially derived amino acids rather than plasma membrane import, for example, through extracellular amino acid deprivation or in cell types with limited surface transporter expression. Our studies have revealed a phenotype best explained by a defect in substrate *export* from mitochondria. This is not necessarily in conflict with other reports mentioned above. A recent preprint reports a matrix-open structure of human Sfxn1 consistent with an alternating-access transport mechanism (Fitzpatrick et al., 2026). Because Sfxn proteins lacks nucleotide-binding and hydrolysis domains, they are unlikely to be ATP-driven primary active transporters. We therefore propose that Sfxn may act as a facilitative transporter, mediating net flux of its substrates with the direction set by cellular context. Our observations suggest that in the germline, solute concentration is higher in the matrix than in the cytoplasm, favoring net export through Sfxn.

The breadth of the amino acid accumulation in mitochondria upon Sfxn depletion raises the question of how much of the phenotype can be explained by a direct loss of Sfxn transport activity versus secondary metabolic consequences. Proposed Sfxn substrates are among those enriched, including serine, proline, threonine, and the branched chain amino acids (BCAAs) leucine and isoleucine, but glutamine showed the largest enrichment by far. Its accumulation could reflect a previously unrecognized Sfxn substrate, a metabolic effect, or both. Additional studies are needed to comprehensively characterize Sfxn substrate specificity across species. Alternatively, because BCAAs are essential and thus not synthesized or interconverted, their accumulation in the matrix could reflect impaired catabolism. Glutamine is also a product of the first step of BCAA processing and thus could inhibit this step by mass action. Distinguishing these possibilities will require reconstitution of Sfxn transport activity in defined systems. In addition, metabolic flux experiments that can resolve the fate of individual amino acids within the mitochondrial compartment under different conditions will be important avenues for future exploration.

A notable feature of the metabolic profile of Sfxn-depleted tissues is how little the total amino acid levels change relative to the striking amino acid sequestration observed within the mitochondrial compartment. Outside of a robust and reproducible elevation of β-alanine, a proposed Sfxn substrate (Block et al., 2026), the global amino acid pool in both oocytes and embryos is largely unchanged. This is consistent with the known expression of plasma membrane amino acid transporters in oocytes, which can supply the cytoplasm with amino acids derived from extracellular sources (Kronja et al., 2014). Such buffering may explain why Sfxn-depleted oocytes give rise to embryos that are largely viable: if the metabolic demands of the oocyte are met through plasma membrane import of metabolites (derived from hemolymph and/or surrounding follicle cells) rather than mitochondrial export, then the sequestration of amino acids in the matrix may impose little selective pressure on embryo survival. However, embryos do require zygotically expressed Sfxn protein, as embryos homozygous mutant for a null allele of *Sfxn1-3* die at the end of embryogenesis (0/55 first-instar larvae were homozygous). Additional metabolomics experiments using timed embryo collections across embryogenesis would identify which metabolites ultimately become limiting.

Together, our findings suggest that metabolite compartmentalization and organelle volume homeostasis are not independent mitochondrial properties but are mechanistically coupled through the same inner-membrane transport systems. In the germline, this coupling is tested when mitochondria must tolerate an abrupt, physiological osmotic shock at egg activation, and the matrix’s pre-existing solute composition determines whether that shock is absorbed or becomes pathological. We suggest that homeostatic metabolite handling to withstand acute physiological transitions may extend well beyond the oocyte to any context in which organelles face rapid, developmentally programmed changes in osmotic environment.

## Author contributions

M.P. and R.L. conceived the project. M.P., L.C., and S.S. performed the RNAi screen and hatch rate analyses. C.G., M.B. and D.M. optimized the EM sample fixation and staining protocol. M.P. and L.C. collected and fixed EM samples. M.B. and D.M. prepared and imaged electron microscopy samples with supervision from A.LJ. M.P. segmented and analyzed EM images. M.P. performed western blot analysis and ddPCR measurements. C.G. generated mitoTag fly lines and M.P. and C.G. validated the lines. M.P. prepared and analyzed metabolomics samples with assistance from S.B. M.P., C.G. and A.A. performed blue native gel assays and in-gel activity assays. M.P. and S.B. performed mitochondrial swelling assays. M.P. performed double knockdown experiments and larval wing disc experiments. M.P., S.B., N.K., and R.L. discussed the Sfxn RNAi phenotype and possible functions. R.L. supervised the work. M.P. wrote the manuscript, and all authors discussed the results and edited the manuscript.

## Acknowledgements

We thank the TRiP at Harvard Medical School (NIH/NIGMS R01-GM084947) for providing transgenic RNAi fly stocks used in this study. We thank the Metabolite Profiling Core Facility at the Whitehead Institute for running metabolomics samples and the Proteomics Core Facility at the Whitehead Institute for preparing and running proteomics samples. TRiP lines were maintained and distributed by the Bloomington Drosophila Stock Center. We are grateful to Matthew Vander Heiden for discussion. We thank Ryan Cummings for guidance on larval wing disc dissections, Agnel Sfeir, Sherilyn Grill, Arjuna Rajakumar, and Mikala Capage for critical reading of the manuscript, and all members of the Lehmann lab for discussions and feedback. This work was supported in part by the Koch Institute Support (core) Grant P30-CA14051 from the National Cancer Institute. We thank the Koch Institute’s Robert A. Swanson (1969) Biotechnology Center for technical support, specifically Peterson (1957) Nanotechnology for Integrative BioEM Core Facility (RRID:SCR_018674) M.P. was supported by NIH/NIGMS F32 GM140745. C.G. was supported by the Swiss National Science Foundation (SNSF) Postdoc Mobility Fund. R.L. acknowledges support from Whitehead Institute Start-up funds.

## Methods

### *D. melanogaster* Strains and fly husbandry

*D. melanogaster* were maintained on cornmeal molasses yeast medium at 25°C using standard procedures. All strains generated and used in this study are listed in the Supplemental Experimental Procedures. DNA constructs for transgenes and insertions were generated by standard methods and injected into D. melanogaster by BestGene.

### Egg hatching rate

To determine egg hatching rate, flies were collected within 24 hours after eclosing and transferred to fresh vials containing cornmeal media to mate for 2-4 days. Flies were then transferred to custom housing overlaying apple juice plates supplemented with yeast paste on the food surface. Each compartment contained 3-7 females and 3-5 males and each condition set up in duplicate. Flies were transferred to new food twice in a 24-hour period. Plates were incubated for 26-36 hours at 25C before counting. Hatch rate analysis reported here was pooled from at least two biological replicates and at least 15 females across all experiments. Hatch rate was determined by dividing the total number of hatched eggs divided by the number of total eggs laid.

### Tissue fixation

We used standard protocols to collect embryos for fixation. Briefly, flies were collected within 3-5 days after eclosing, sorted, and transferred to a custom cage overlaid on an apple juice place supplemented with yeast paste. After flies acclimated to the cage for 2 days, the apple juice plates were swapped, and flies were allowed to lay eggs on the plates for the desired duration. To fix, extra yeast paste was removed, and a fresh 50% bleach solution was applied to the plate. Embryos were gently dislodged from the agar using a cell scraper and swirled gently in the bleach solution until 50-80% of the dorsal appendages had dissolved (about 4 minutes). Embryos were collected by straining through a fine mesh and washed thoroughly with water. Embryos were transferred using a fine paint brush to a glass vial containing a 1:1 mixture of 4% PFA in 1x PBS and saturated heptane (shaken thoroughly with fixative solution before transferring to the glass vial). Samples were shaken on a rotary shaker at 120 rpm for 20 minutes. The aqueous (lower) fraction was removed and 10 ml methanol was added. Vials were immediately shaken rapidly for 30 seconds to remove the vitelline membrane. Embryos were transferred to an Eppendorf tube, washed 3x with fresh methanol over a 10-minute period, and stored at −20 °C until needed.

To collect ovary and oocyte samples for fixation, we applied a similar protocol. Flies were anesthetized using FlyNap and transferred to a drop of PBS-Tw. Ovaries were removed from up to 10 flies over a 10-minute period using fine #55 Dumont tweezers and transferred to an Eppendorf tube containing PBS-Tw. Egg chambers were dissociated from the ovary by pipetting rapidly using a p200 pipette tip. The buffer was exchanged for 4% PFA in 1X PBS and incubated for 30 minutes. Fixative was removed and oocytes washed twice with PBS-Tx. Samples were either stored at 4C for a few days until needed or transferred to vectashield + DAPI and allowed to incubate at least overnight before imaging.

### Electron microscopy

Ovaries and oocytes for electron microscopy were prepared for fixation as described above, but placed in a fixation buffer of 2% paraformaldehyde and 2.5% glutaraldehyde in PBS for 1 hour. Fixative was removed and samples washed 3 times with 0.1M sodium cacodylate buffer, and incubated in 1% Osmium tetroxide and 1.25% Potassium ferrocyanide at 4C for one hour. Samples were then rinsed 3 times in 50mM maleate buffer and incubated overnight in 2% uranyl acetate in maleate buffer. Samples were then dehydrated through a graded series of ethanol, transitioned through propylene oxide, and infiltrated with Eponate 812 resin. Sample blocks were polymerized at 60C for at least 48 hours.

Blocks were trimmed and sectioned on a Leica UC7 ultramicrotome. 60nm ultrathin sections were cut using a DiATOME ultra 45 diamond knife. Sections were collected on piloform coated copper 2×1mm slot grids.

Imaging was performed on a Zeiss Crossbeam 540 Scanning Electron Microscope (SEM) using an insertable STEM detector and operating at 30kV and 200pA. Image mosaics were acquired using the Zeiss ATLAS 5 software at 4nm pixel density and 6.4µs dwell time. Mosaics from an individual embryo or ovary were stitched into a single montage, and exported as TIFF files for subsequent analysis.

### ddPCR mtDNA copy number measurements

ddPCR reactions were carried out according to the manufacturer’s (Bio-Rad) protocol. ddPCR droplets were generated from samples using QX200 Droplet Generator (Bio-Rad) and underwent complete PCR cycling on a C100 deep-well thermocycler (Bio-Rad). Droplet fluorescence was read using the QX200 Droplet Reader (Bio-Rad). Sample copy number was determined using Quantasoft software (Bio-Rad).

### Western blot

To determine the relative amounts of protein expression in control and Sfxn knockdown embryos, we collected 0-2 hr old embryos and dechorionated in 50% bleach for 2-4 minutes. We handpicked 20 embryos per sample and transferred them to an Eppendorf tube using a fine paintbrush or needle. Samples were lysed immediately or stored in −20 °C until needed. To lyse, we added 50 μl of 1x Laemmli sample buffer containing BME to each sample and homogenize with a pestle for 15 seconds. To reduce foaming, samples were spun on a tabletop centrifuge for 3 minutes. Samples were then heated in a heat block for 1 minute at 95 °C then cooled to room temperature before loading in a gel. 5 µl of each sample was run on a 4–15% Mini-PROTEAN® TGX™ Precast Protein Gels, 10-well, 50 µl #4561084, then transferred to nitrocellulose membrane using the BioRad TransBlot Turbo system for blotting. The membrane was incubated in BioRad blocking buffer for 30 minutes before transferring to primary antibody solution. Li-Cor secondary antibodies were used at 1:20,000 dilution. Blots were exposed on a LiCor Odyssey CLx scanner.

### Metabolomics

For whole tissue lysates, stage 14 oocytes were collected from ∼100 isogenic females expressing either *sfxn* shRNA or nontargeting control shRNA. The same number of stage 14 oocytes were manually dissected for each sample and analyzed in triplicate (n = 200 x 3 or n = 100 x 3). For embryo samples, embryos were collected from >200 females, pooled, then split into triplicate samples (up to 300 embryos per triplicate sample). Samples were flash-frozen in liquid nitrogen and stored at −80°C. To extract metabolites, an 80/20 mixture of cold 80% methanol supplemented with 20 ^13^C-labeled amino acid standards (see Supp Table 3) was added to the samples along with ∼30 ceramic beads of Lysing Matrix E from MP Biologicals. Samples were homogenized using a Qiagen TissueLyser II bead mill by shaking for a minimum of 2 minutes and 30 seconds at 30 Hz at room temperature and visually inspected to confirm homogenization. If needed, samples were subjected to a second round of shaking under the same conditions. The homogenate was transferred to a fresh 1.5 mL Eppendorf tube, and the beads were rinsed with fresh extraction buffer and combined with the homogenate. Samples were spun at max speed on a tabletop centrifuge for 10 mins at 4°C. The supernatant was carefully removed and transferred to a fresh 1.5 mL Eppendorf tube. Samples were dried at room temperature using a flow of nitrogen, then resuspended in LSMS-grade water, spun down again to pellet any remaining precipitate, and then samples were analyzed.

Metabolite profiling was conducted on a QExactive bench top orbitrap mass spectrometer equipped with an Ion Max source and a HESI II probe, which was coupled to a Dionex UltiMate 3000 HPLC system (Thermo Fisher Scientific, San Jose, CA). Samples were injected onto a SeQuant® ZIC®-pHILIC 150 x 2.1 mm analytical column equipped with a 2.1 x 20 mm guard column (both 5 um particle size; EMD Millipore). Buffer A was 20 mM ammonium carbonate, 0.1% ammonium hydroxide; Buffer B was acetonitrile. The column oven and autosampler tray were held at 25⁰C and 4⁰C, respectively. The chromatographic gradient was run at a flow rate of 0.150 mL/min as follows: 0-20 min: linear gradient from 80-20% B; 20-20.5 min: linear gradient from 20-80% B; 20.5-28 min: hold at 80% B. The mass spectrometer was operated in full-scan, polarity-switching mode, with the spray voltage set to 3.0 kV, the heated capillary held at 275⁰C, and the HESI probe held at 350⁰C. The sheath gas flow was set to 40 units, the auxiliary gas flow was set to 15 units, and the sweep gas flow was set to 1 unit. MS data acquisition was performed in a range of *m/z* = 70–1000, with the resolution set at 70,000, the AGC target at 1×10^6^, and the maximum injection time at 20 msec.

Relative quantification of metabolites was carried out using XCalibur 2.2 software (Thermo Fisher Scientific) and/or Skyline (Adams et al., 2020) referencing an in-house library of chemical standards for initial peak picking. The total peak area of stable isotope labeled internal standards in each injected sample was used to confirm reproducibility in sample extraction/injection volume. We manually inspected and verified peak integration profiles, adjusting when necessary. Metabolites with a water-blank signal reproducibly exceeding 50% of the biological median (detected in ≥2/4 blank injections, area ≥50,000) were excluded from quantitative analysis due to analytical background. Peak areas for each metabolite were normalized to a per-sample extraction and injection quality factor derived from stable isotope labeled internal standards. The following sixteen isotopically labeled amino acid standards were used: alanine-13C3-15N, arginine-^13^C6-^15^N4, aspartate-^13^C6-^15^N4, glutamate-^13^C5-^15^N, glycine-^13^C2-^15^N, histidine-^13^C6-^15^N3, Isoleucine-^13^C6-^15^N, leucine-^13^C6-^15^N, lysine-^13^C6-^15^N2, methionine-^13^C5-^15^N, phenylalanine-^13^C9-^15^N, proline-^13^C5-^15^N, serine-^13^C3-^15^N, threonine-^13^C4-^15^N, tyrosine-^13^C9-^15^N, valine-^13^C5-^15^N. These standards were spiked at equal concentration into the extraction solvent prior to metabolite extraction, such that their recovery reflects both extraction efficiency and injection volume for each sample.

Data was analyzed by Metaboanalyst (Pang et al., 2024) to perform fold-change analysis and determine significance. Graphs were plotted in Prism.

### Mitochondrial swelling assays

Mitochondrial isolation and swelling assays were carried out using a previously described protocol, modified for drosophila tissue (Block et al., 2026). Briefly, ∼2000 embryos were collected within 2 hours after egg laying, rinsed with cold PBS. Embryos were transferred to a dounce homogenizer and cold isolation buffer (250 mM sucrose, 10 mM Tris pH 7.5, 10 mM KCl, 1.5 mM MgCl_2_, 1 mM potassium EDTA) was added. Embryos were lysed using 30 strokes + twisting of a dounce homogenizer with a tight fit pestle. Nuclei and cell debris were removed with two 10-minute spins at 1300 x g at 4C. Mitochondria were isolated by differential centrifugation by spinning for 10 mins at 10,000 x g at 4C. The pelleted mitochondria were washed with cold isolation buffer and spun again. The mitochondria pellet was resuspended in room-temperature swelling buffer (20 mM Tris pH 7.5, 0.5 mM potassium EDTA, 10 μM rotenone and adjusted to 2 mg/ml. The suspension was incubated at room temperature for 30 minutes before swelling was initiated by adding an equal volume of room temperature swelling buffer containing 500 mM substrate (250 mM final concentration). Swelling was monitored by measuring absorbance at 520 nm using a microplate reader, and swelling rates were determined as the absolute value of the change in absorbance at 520 nm from 0 to 90 seconds. Swelling rates were normalized to simultaneously swollen WT mitochondria unless otherwise stated.

For glutamine uptake, which is less soluble than alanine, mitochondria in isolation buffer were incubated at room temperature for 30 minutes and subsequently swollen with swelling buffer containing 200 mM glutamine and 50 mM sucrose.

---

**Supplemental Figure 1.**
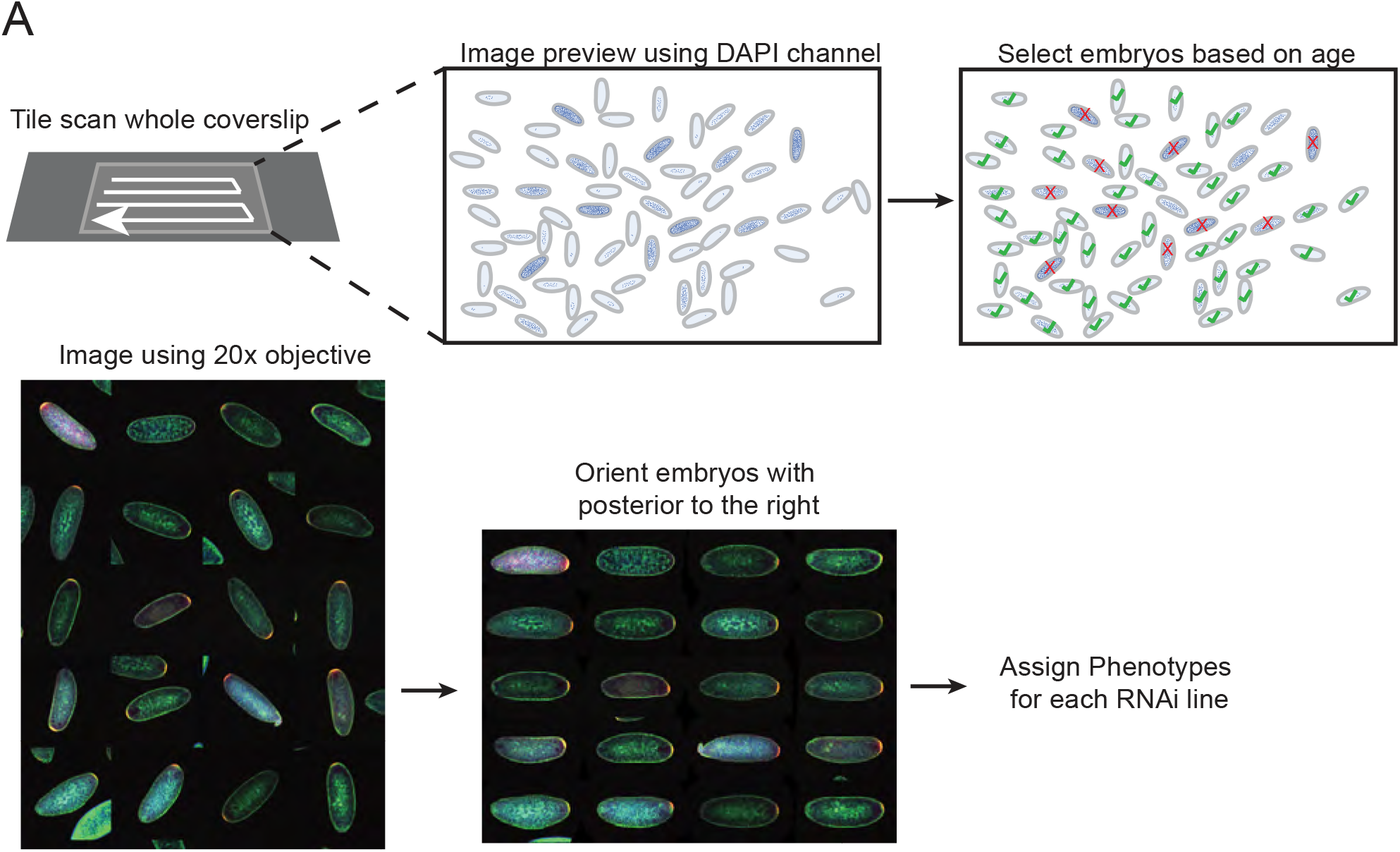
A. RNAi screen imaging workflow. Fixed and immunostained embryos aged 0-2 hr old were mounted on a slide. A tile-stitched overview image was generated using DNA channel (DAPI stain) to capture all embryos on a coverslip. Embryos in nuclear cycle 1-9 were selected for 20x imaging in all three channels to visualize mitochondria, germplasm, and DNA. Phenotypic categories were determined by visual assessment of embryo montages.

**Supplemental Figure 2.**
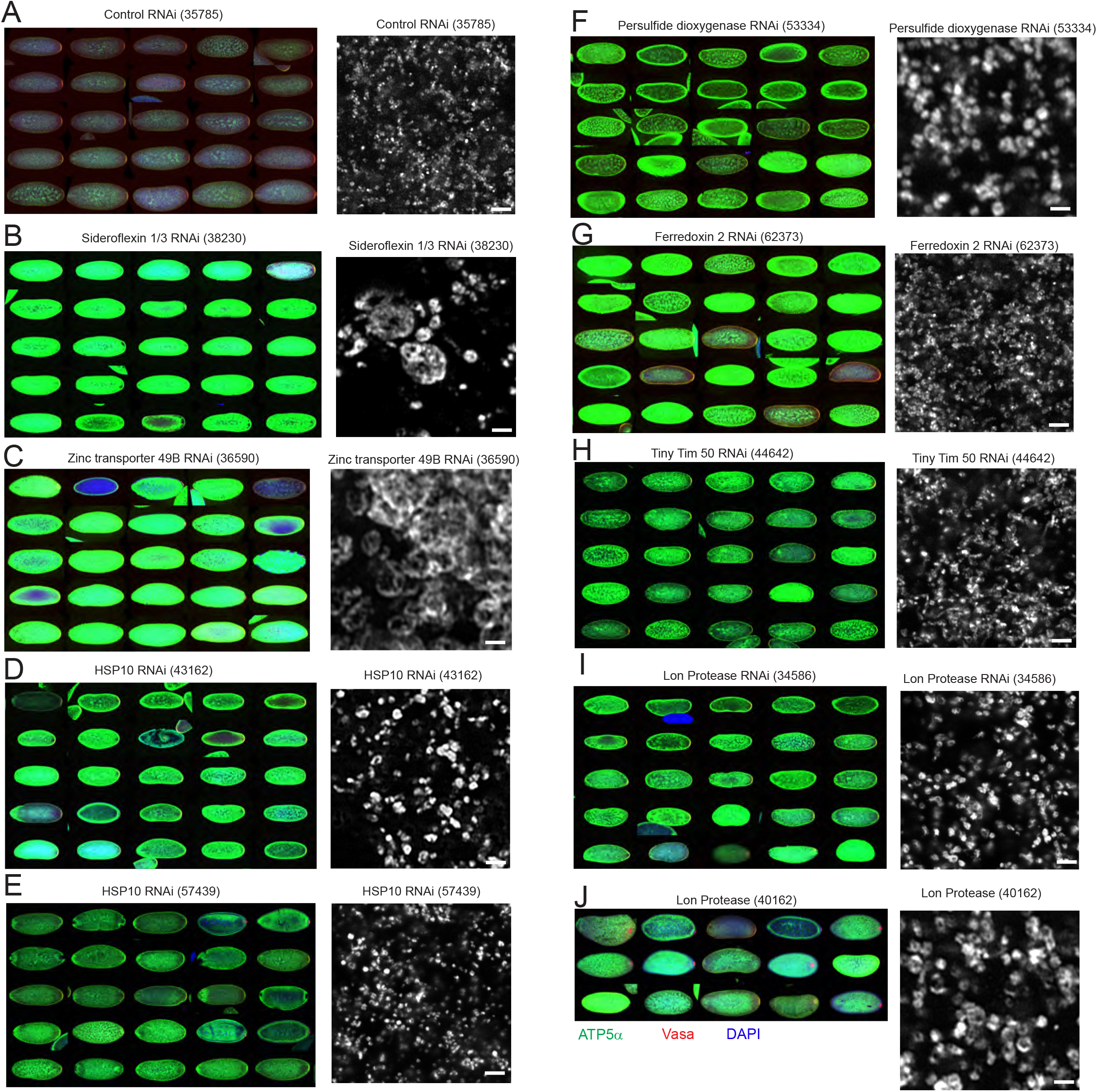
A-J. Laser scanning confocal images of embryos from RNAi lines exhibiting the Bright phenotype. Left: Montages of 20x images immunostained for mitochondria (anti-ATP5α, green), germplasm (anti-Vasa, red) and DNA (DAPI, blue). Right: 63x airyscan images of mitochondria (anti-ATP5α). Scale bar: 2 µm.

**Supplemental Figure 3.**
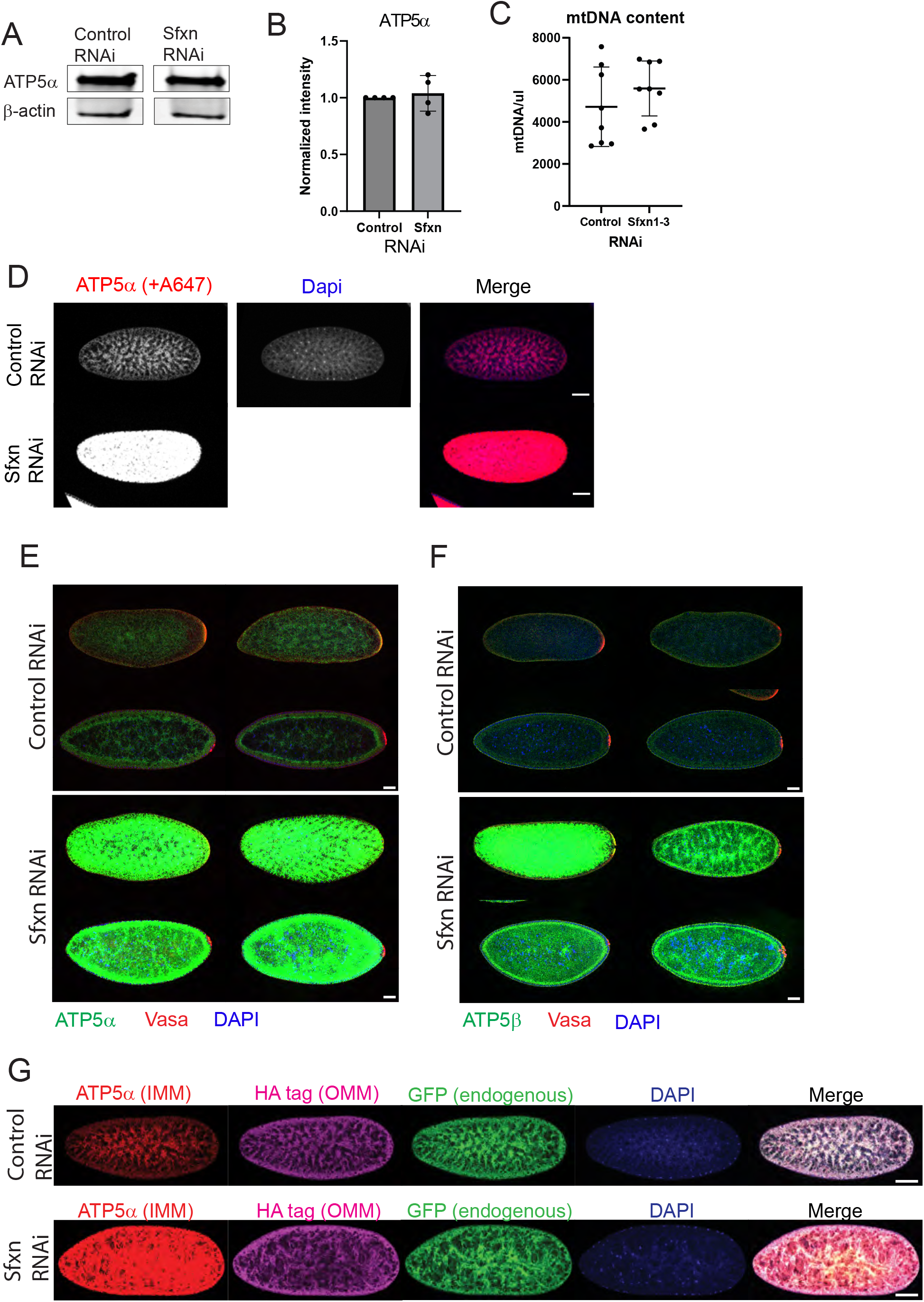
A. Example western blot showing ATP5α levels in 0-2 hr old embryos from control RNAi and Sfxn RNAi conditions. β-actin is shown as a loading control. B. Quantification of western blots probed with ATP5α antibody from n = 3 independent experiments. C. Droplet digital PCR (ddPCR) analysis of mtDNA content per microliter of prepared embryo lysate. Each experiment consisted of pooled lysates from 10 embryos aged 0-2 hr old. D. 20x laser scanning confocal images of immunostained embryos. Top: control embryos. Bottom: Sfxn RNAi embryos. Left: Single-channel image of mitochondria channel, visualized using a mouse anti-ATP5α primary antibody and an anti-mouse 647 secondary antibody. All other images in this manuscript use an anti-mouse 488 secondary antibody. Left: Single-channel image showing ATP5α staining. Middle: Single channel images showing DAPI staining. Right: two-color overlay image showing ATP5α in red and DAPI in blue. E-F. 20x laser scanning confocal image montages. Top: Control embryos. Bottom: Sfxn RNAi embryos. The same microscope and display settings are applied across all images. Three-channel overlay images show germplasm in red (anti-Vasa), DNA in blue (DAPI), and mitochondria in green, stained with either (B) anti-ATP5α or (C) ant-ATP5β. Scale bar: 50 µm. G. 20x laser scanning confocal image montages. Embryos are derived from mothers expressing ‘MitoTag’ (GFP and HA on outer mitochondrial membrane) and shRNA to a nontargeting control sequence (top) or *Sfxn1-3* (bottom). Images were collected using the same microscope settings across experiments. The same display settings for each channel were applied to control and Sfxn RNAi images. Anti-ATP5α (in the mitochondrial matrix) is shown in red, anti-HA (on the outer mitochondrial membrane) is shown in magenta, endogenous GFP fluorescence (on the outer mitochondrial membrane) is shown in green. DNA marked by DAPI is shown in blue. Scale bar: 50 µm.

**Supplemental Figure 4.**
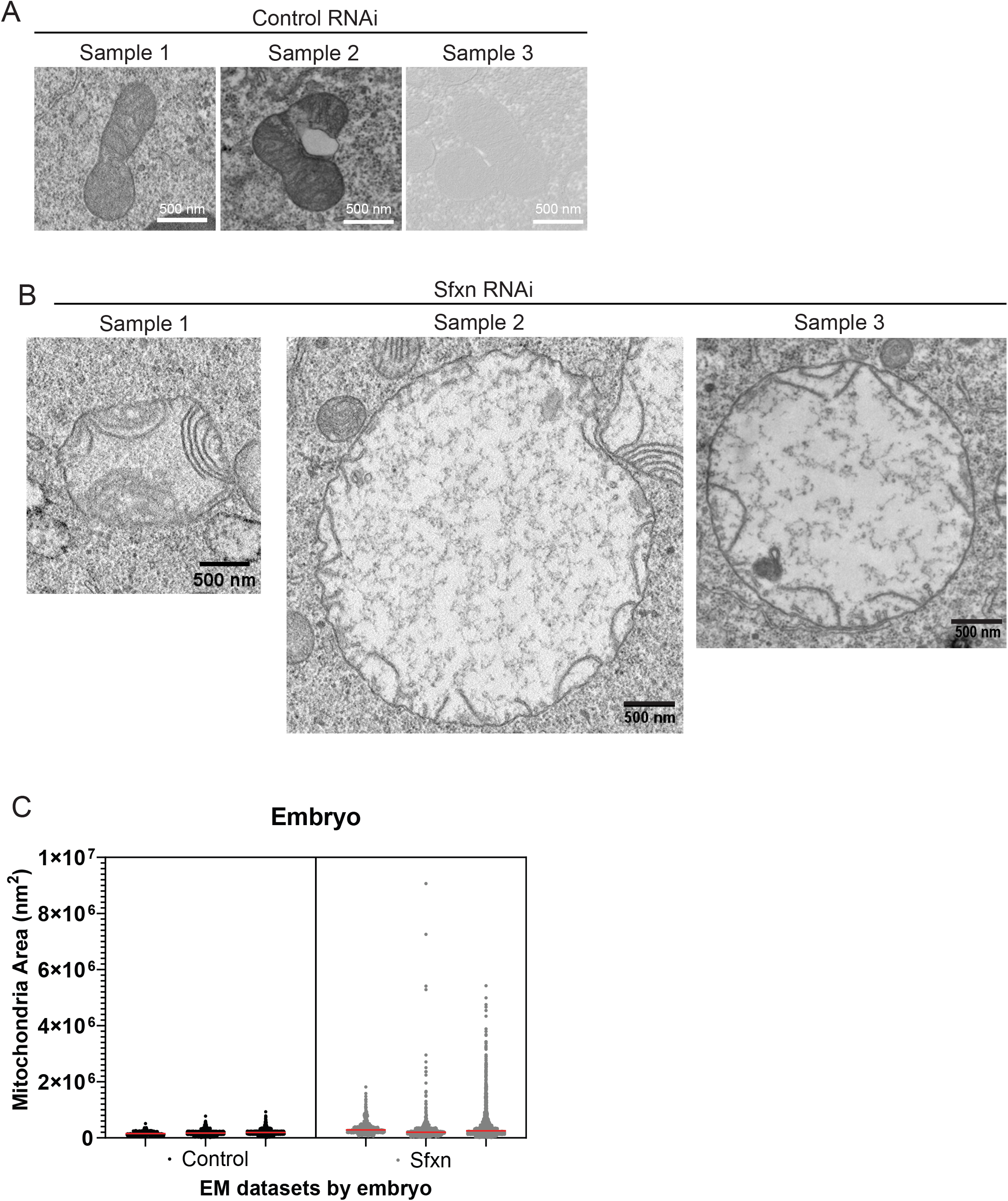
A-B. Images of the largest mitochondrion in each (A) control or (B) Sfxn RNAi embryo in EM datasets. C. Violin plot of mitochondrial area in EM data, separated by dataset. n = 3 embryos per condition.

**Supplemental Figure 5.**
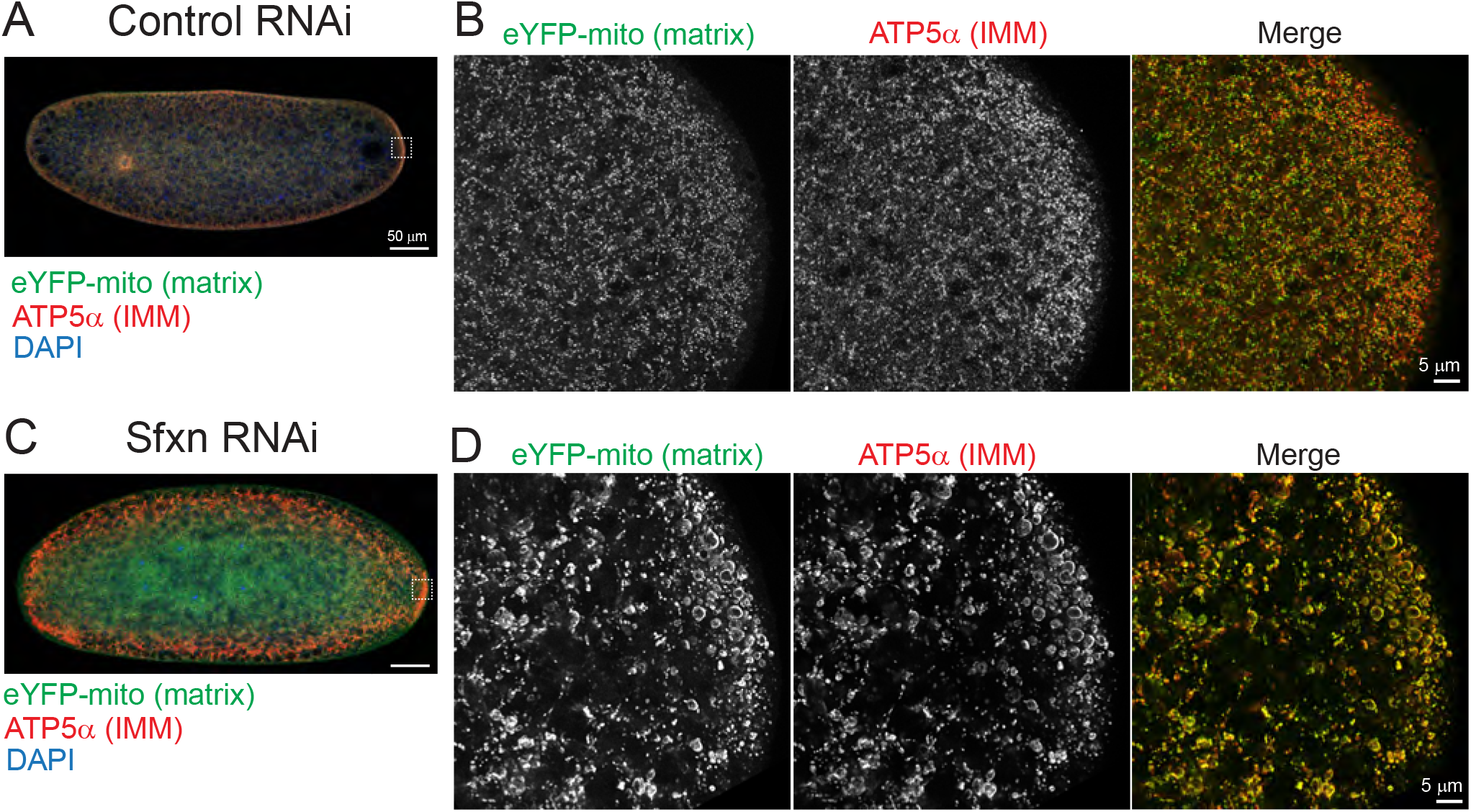
A-D. Colocalization of endogenously expressed eYFP-mito construct, which localizes to the mitochondrial matrix, with anti-ATP5α immunostain, which targets a protein in the inner mitochondrial membrane (IMM). An example embryo from (A-B) control or (C-D) Sfxn RNAi conditions.

**Supplemental Figure 6.**
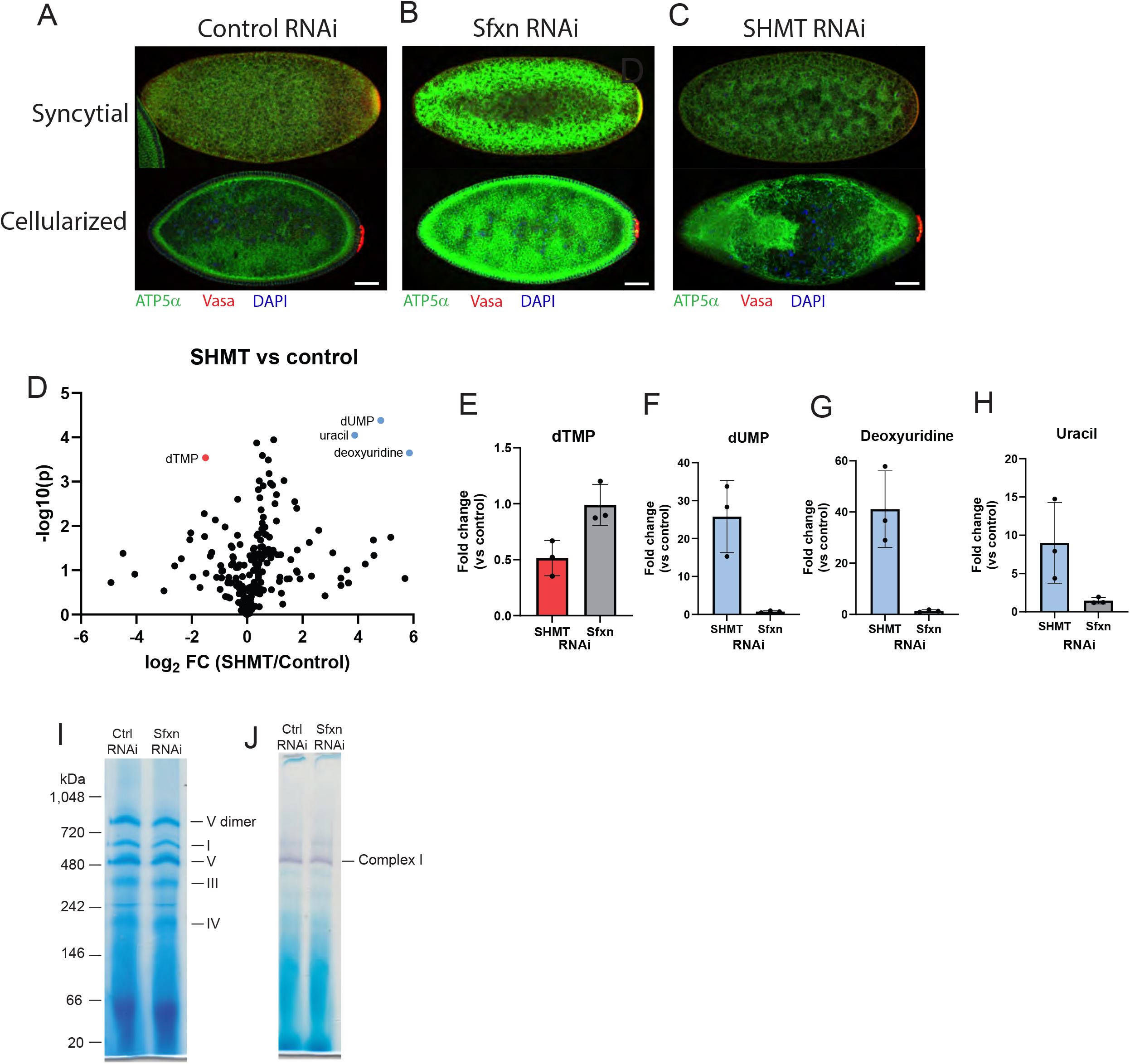
A-C. Example immunofluorescence images from syncytial (stage 2) (top) and cellularized (stage 4) (bottom) stages of embryos expressing shRNA to (A) control, (B) *Sfxn1-3*, or (C) *Shmt*. Green: ATP5α; red: Vasa; DNA (DAPI). Scale bar: 50 µm. D. Volcano plot from an example metabolomics experiment. Results from whole-tissue metabolomics from SHMT-depleted embryos compared to control embryos. E-H. Bar charts showing fold-change values for indicated metabolites in embryos expressing shRNA to SHMT or Sfxn relative to control. Data are from three independent experiments. Dots represent fold-changes values averaged from 3 technical replicates per experiment. I. Blue native gel showing assembled electron transport chain (ETC) complexes. Lysates were prepared from (lane 1) control or (lane 2) Sfxn RNAi embryos. Bands corresponding to different ETC complexes are indicated. J. In-gel activity assay for complex I function. Lysates were prepared from (lane 1) control or (lane 2) Sfxn RNAi embryos.

**Supplemental Figure 7.**
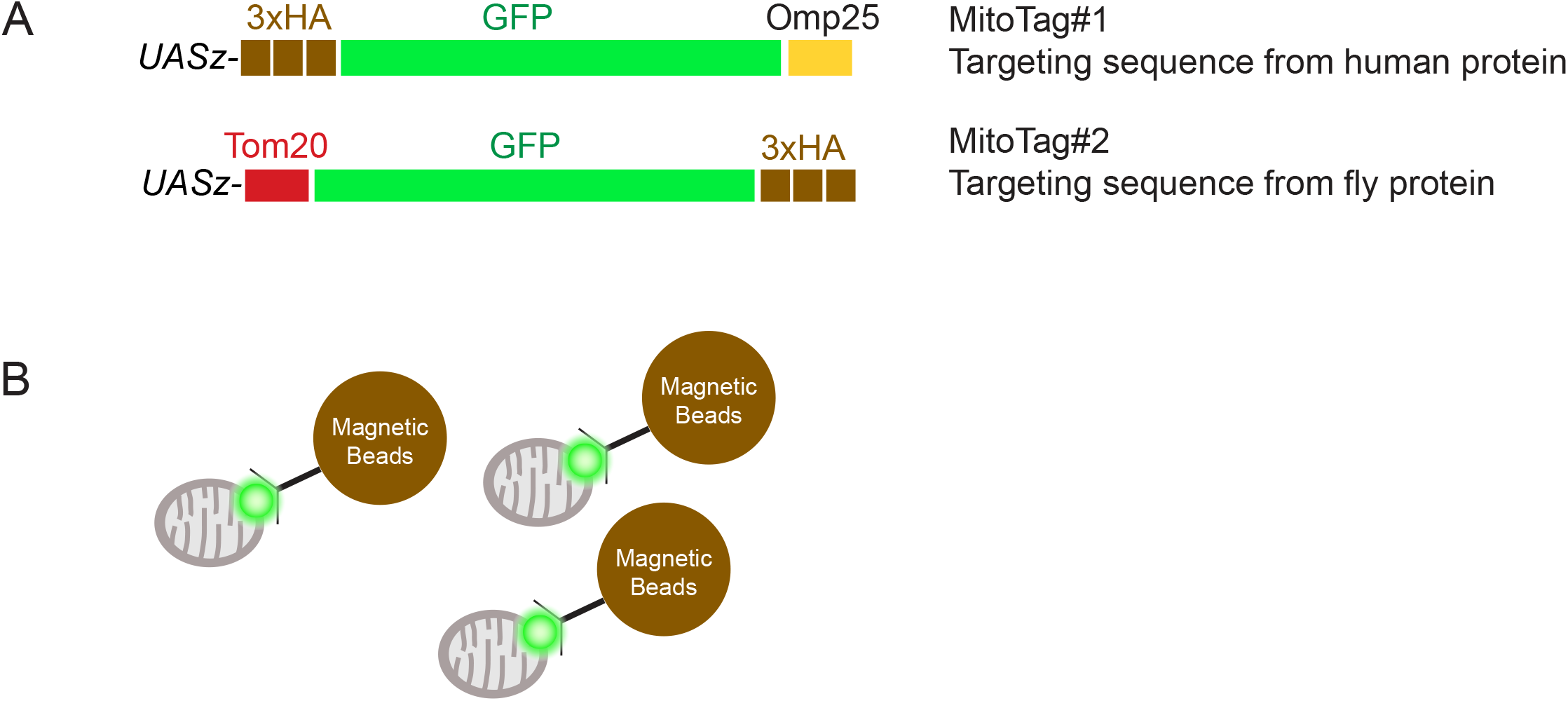
A. Schematic of *UASz*-mitoTag construct design with corresponding description. B. Cartoon of mitoTag-based immunopurification of mitochondria.

**Supplemental Figure 8.**
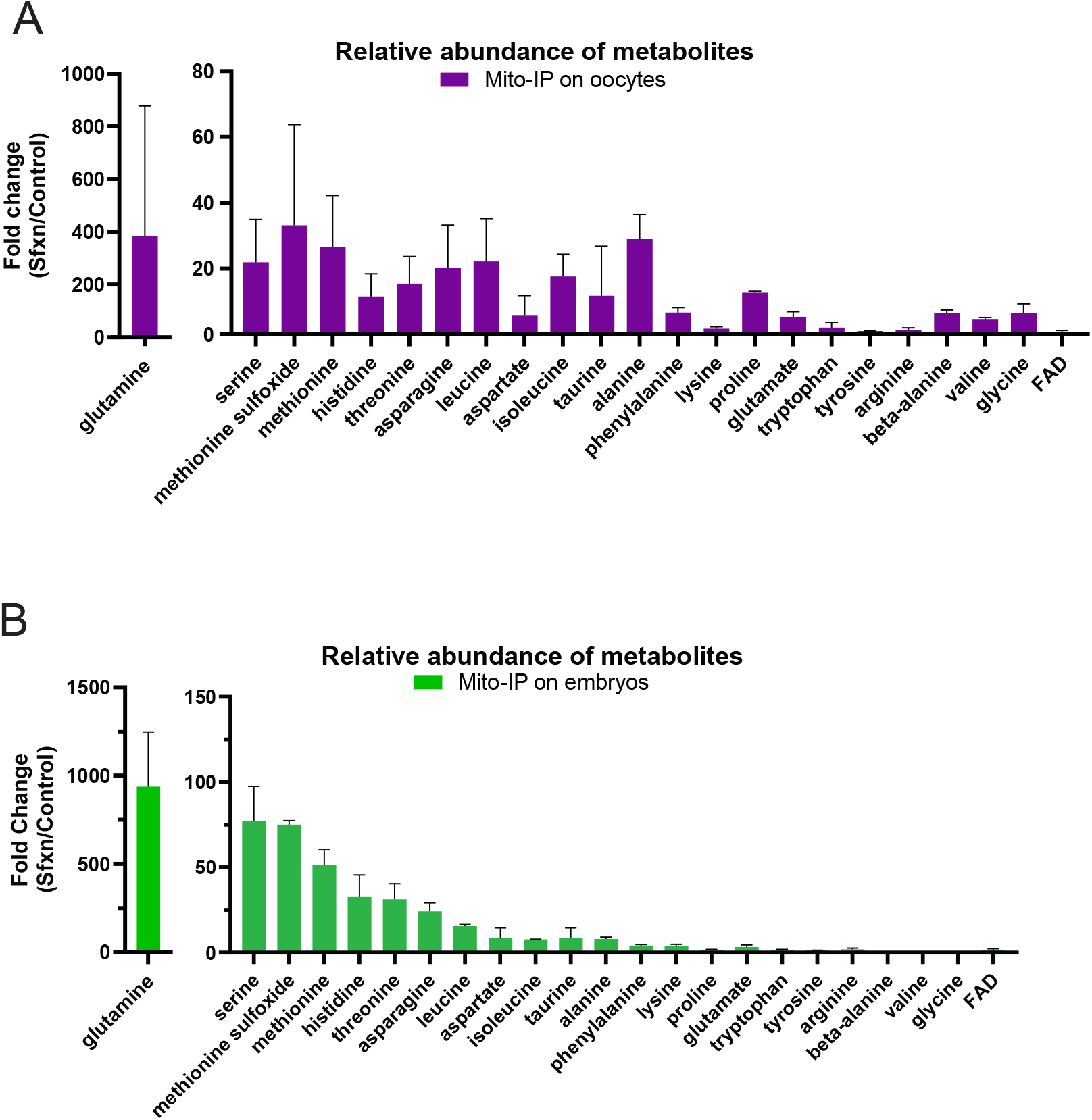
A-B. Metabolomics on immunopurified mitochondria. Data shown are from Fig 4 F and H, with Y axes modified. (A) Data from Figure 4F. (B) Data from Figure 4H. Bar charts show relative abundance of indicated amino acids and metabolites. All measurements are metabolite abundance in Sfxn RNAi samples relative to control RNAi samples. Mean +/− S.D. Embryo samples are from syncytial stage (aged 0-2 hr old). Oocyte samples are from mature (stage 14) oocytes.

**Supplemental Table 1.**
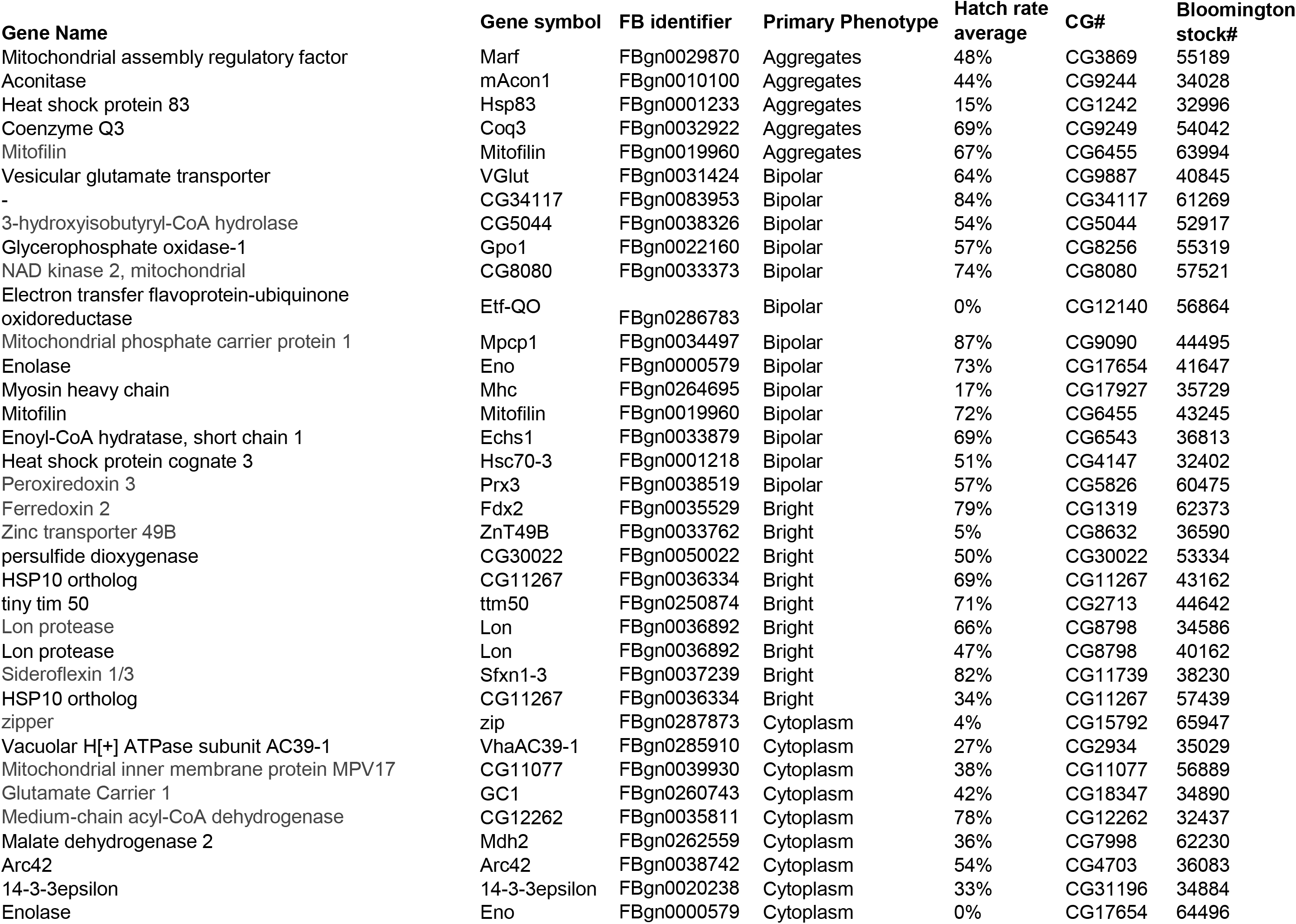
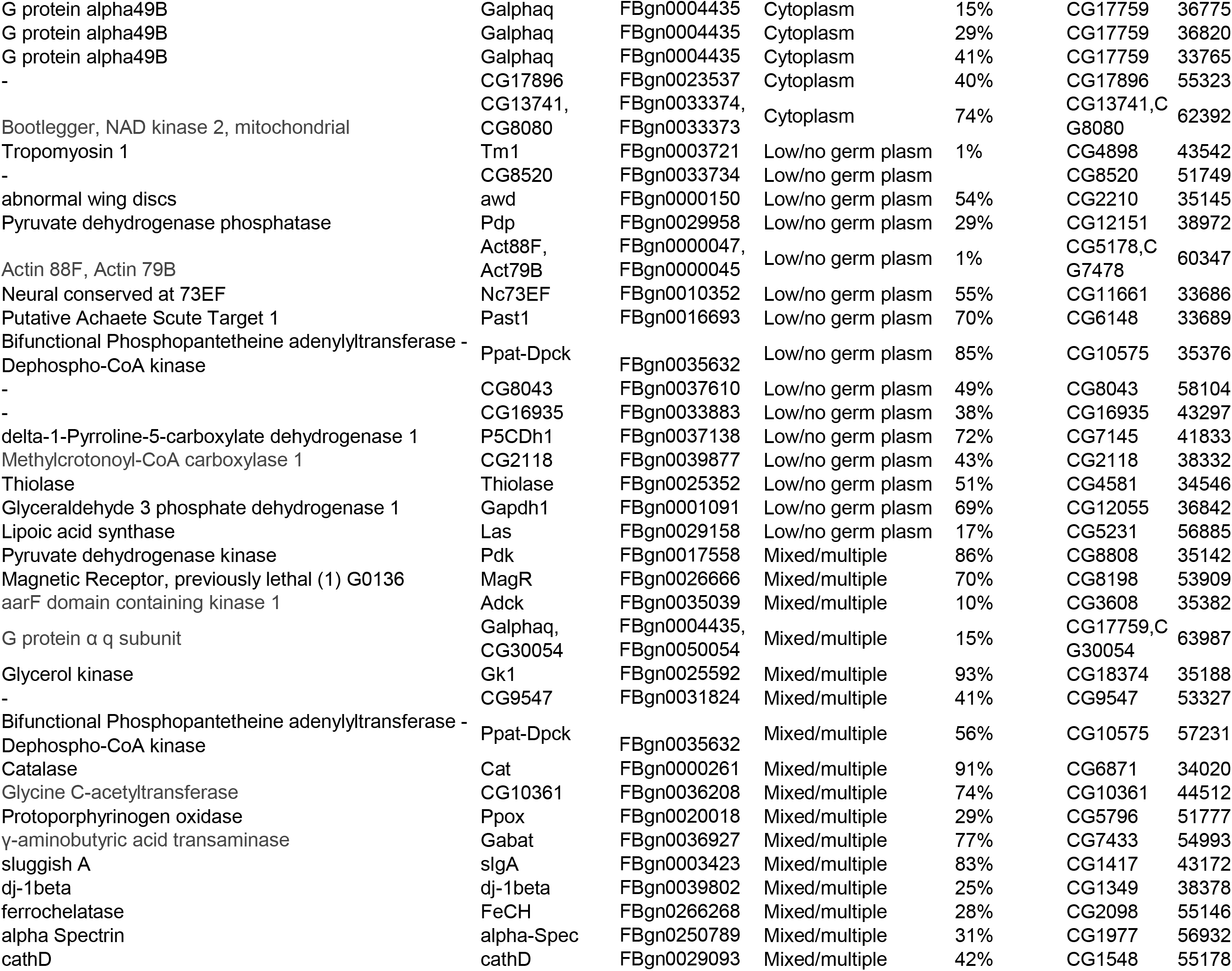
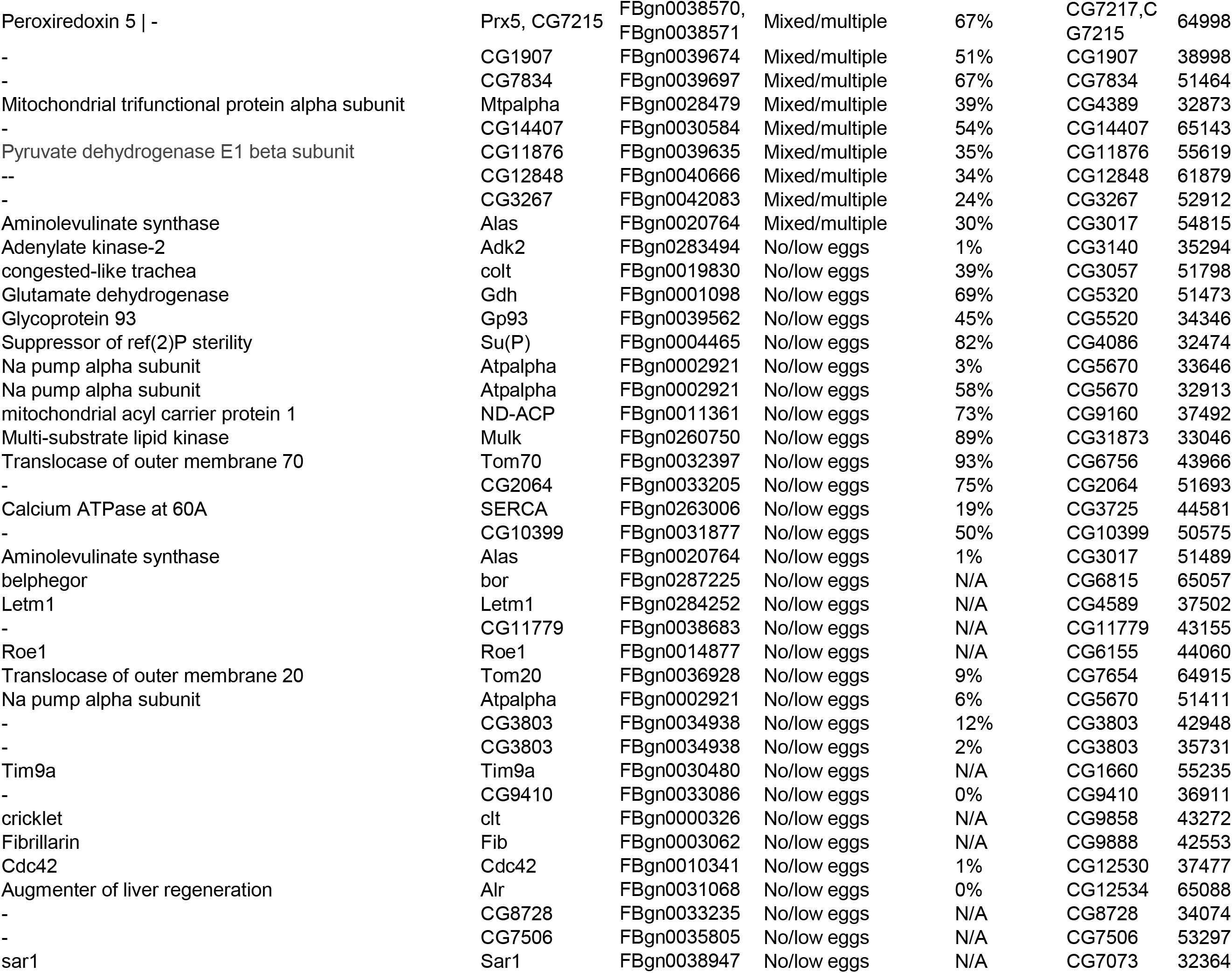
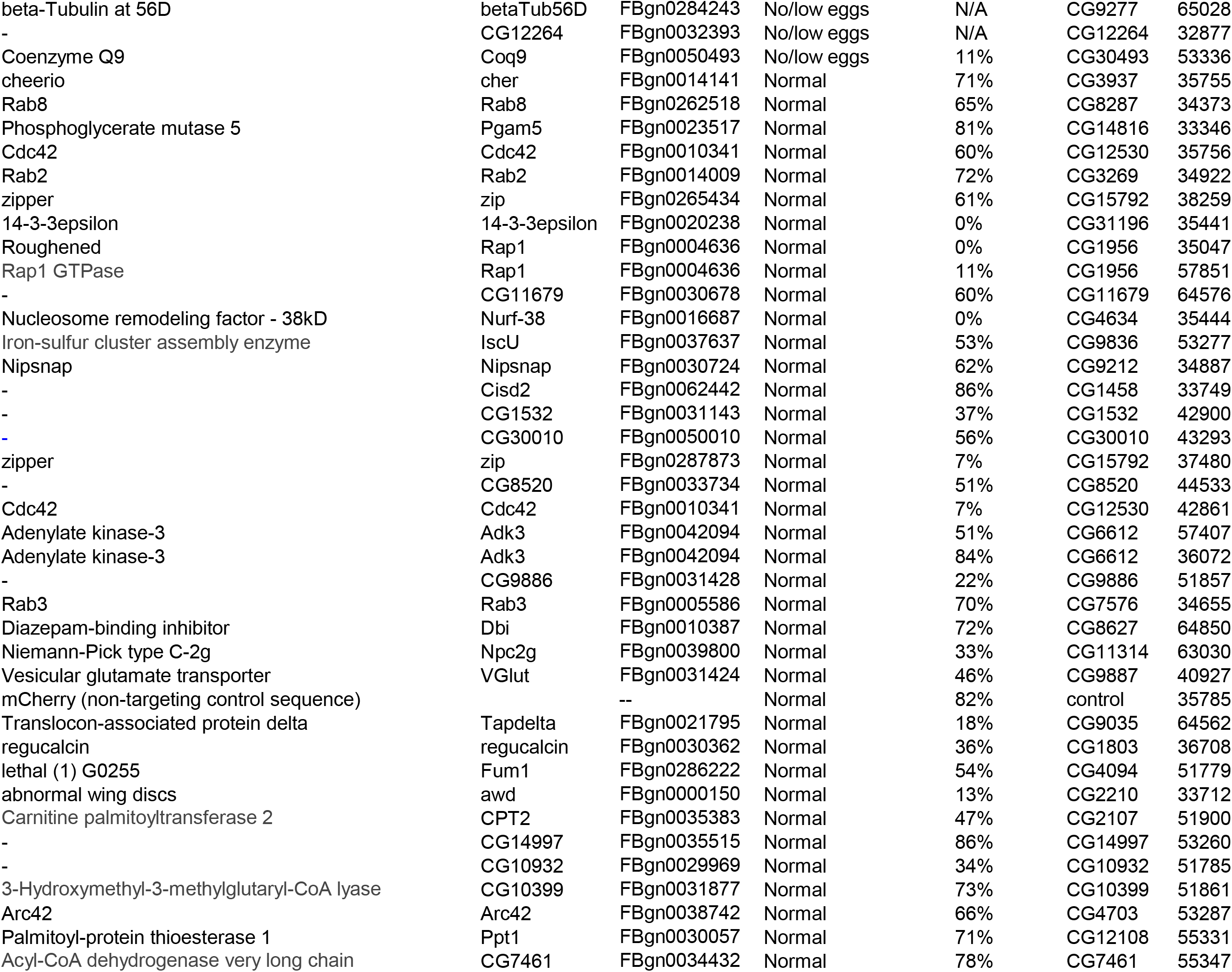
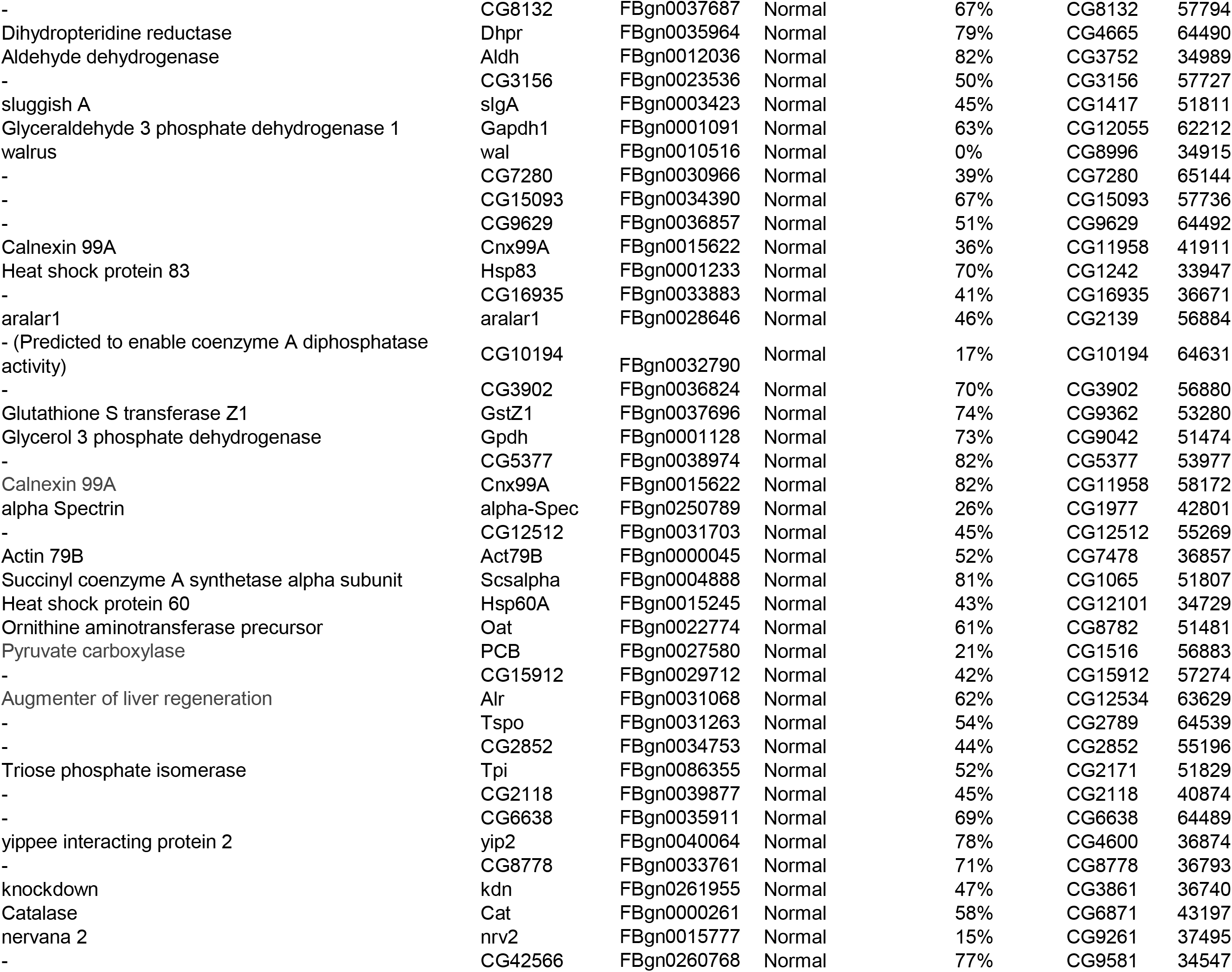
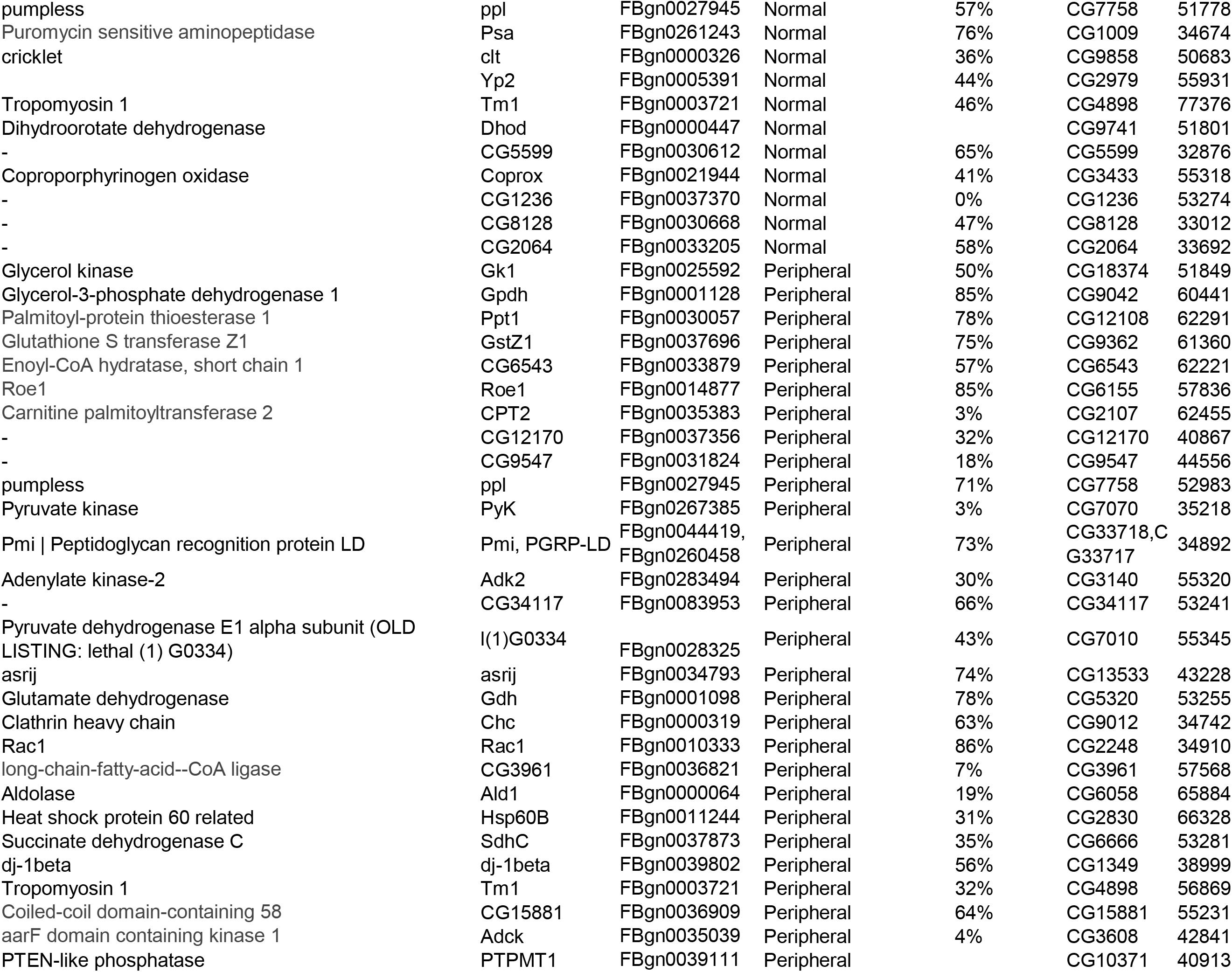
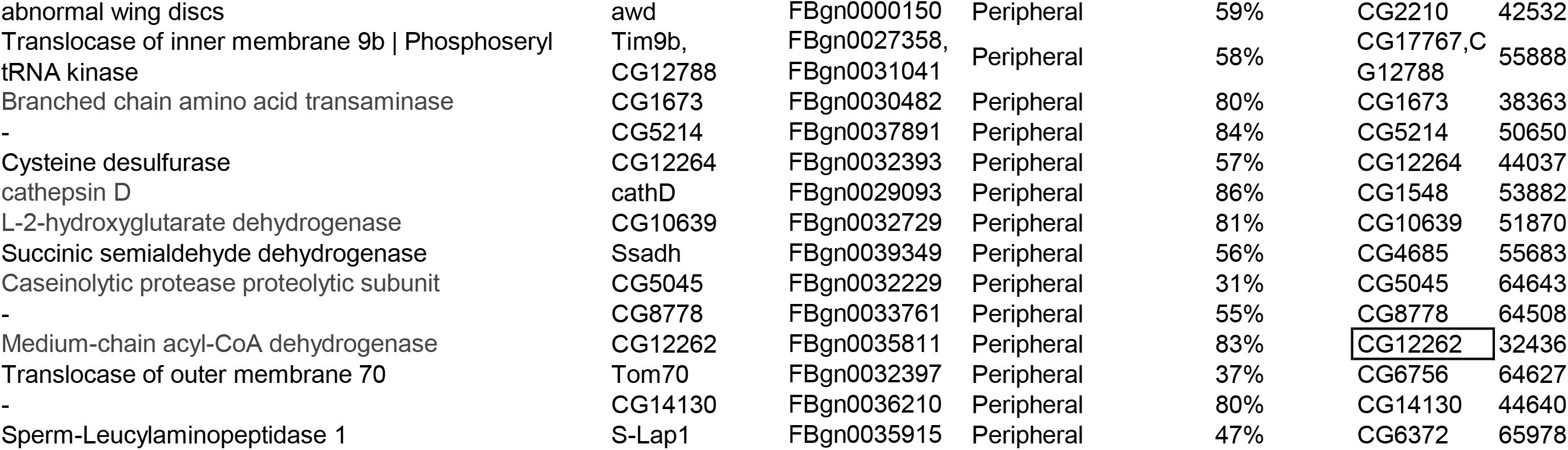
Results from RNAi screen. For each TRiP line, table shows phenotypic class assignment, gene targeted, and hatch rate along with CG and FlyBase identifier.

**Supplemental Table 2.** List of polar metabolites in targeted metabolomics experiments. For each metabolite, Label Type, Molecular Formula, Precursor Adduct, Precursor m/z, and Explicit Retention Time are shown.

| Molecule | Label Type | Molecular Formula | Precursor Adduct | Precursor m/z | Explicit Retention Time |
| --- | --- | --- | --- | --- | --- |
| 1-methyladenosine | light | C11H15N5O4 | [M+H] | 282.1197 | 6.50 |
| 1-methylnicotinamide | light | C7H9N2O | [M+H] | 137.0709 | 12.60 |
| 2,3-BPG | light | C3H8O10P2 | [M-H] | 264.9520 | 12.40 |
| 2-aminoadipate | light | C6H11NO4 | [M+H] | 162.0761 | 10.30 |
| 2-Hydroxybutyric_acid (AHB) | light | C4H8O3 | [M-H] | 103.0401 | 4.53 |
| 2-hydroxyglutarate | light | C5H8O5 | [M-H] | 147.0299 | 10.77 |
| 2-oxoadipate | light | C6H8O5 | [M-H] | 159.0299 | 9.80 |
| 3-hydroxybutyrate | light | C4H8O3 | [M-H] | 103.0401 | 5.50 |
| 3-hydroxyisobutyric_acid | light | C4H8O3 | [M-H] | 103.0401 | 6.14 |
| 3-methylglutaryl carnitine | light | C13H23NO6 | [M+H] | 290.1598 | 8.1 |
| 3-phosphoglycerate | light | C3H7O7P | [M-H] | 184.9857 | 11.70 |
| 3-ureidopropionate | light | C4H8N2O3 | [M+H] | 133.0608 | 8.00 |
| 4-Guanidinobutanoic acid | light | C5H11N3O2 | [M+H] | 146.0924 | 9.50 |
| 4-Hydroxybutyric acid | light | C4H8O3 | [M-H] | 103.0401 | 6.52 |
| 5-Hydroxyindole-3-Acetate | light | C10H9NO3 | [M+H] | 192.0655 | 8.50 |
| 6-phosphogluconate | light | C6H13O10P | [M-H] | 275.0174 | 12.30 |
| 7-methylguanosine | light | C11H16N5O5 | [M+H] | 298.1146 | 7.58 |
| acetoacetate | light | C4H6O3 | [M-H] | 101.0244 | 4.60 |
| Acetyl CoA | light | C23H38N7O17P3S | [M+H] | 810.1330 | 8.50 |
| acetylcarnitine | light | C9H17NO4 | [M+H] | 204.1230 | 5.85 |
| acetylserine | light | C5H9NO4 | [M-H] | 146.0459 | 6.80 |
| adenine | light | C5H5N5 | [M+H] | 136.0618 | 4.50 |
| adenosine | light | C10H13N5O4 | [M+H] | 268.1040 | 4.10 |
| ADP | light | C10H15N5O10P2 | [M-H] | 426.0221 | 10.4 |
| ADP-ribose | light | C15H23N5O14P2 | [M-H] | 558.0644 | 10.3 |
| AICAR | light | C9H15N4O8P | [M-H] | 337.0555 | 10.5 |
| alanine | light | C3H7NO2 | [M+H] | 90.0550 | 9.22 |
| alanine-13C3-15N | heavy | C'3H7N'O2 | [M+H] | 94.0621 | 9.22 |
| allantoin | light | C4H6N4O3 | [M-H] | 157.0367 | 8.5 |
| alpha-aminobutyric acid | light | C4H9NO2 | [M+H] | 104.0706 | 8.27 |
| Alpha-Ketoglutarate (aKG) | light | C5H6O5 | [M-H] | 145.0143 | 10.92 |
| Alpha-Lipoic Acid | light | C8H14O2S2 | [M-H] | 205.0362 | 2.2 |
| AMP | light | C10H14N5O7P | [M-H] | 346.0558 | 9.44 |
| anserine | light | C10H16N4O3 | [M+H] | 241.1295 | 9 |
| arginine | light | C6H14N4O2 | [M+H] | 175.1190 | 13.85 |
| arginine-13C6,15N4 | heavy | C'6H14N'4O2 | [M+H] | 185.1273 | 13.85 |
| argininosuccinate | light | C10H18N4O6 | [M+H] | 291.1299 | 11.5 |
| asparagine | light | C4H8N2O3 | [M+H] | 133.0608 | 9.6 |
| aspartate | light | C4H7NO4 | [M-H] | 132.0302 | 10.3 |
| aspartate-13C6-15N4 | heavy | C'4H7N'O4 | [M-H] | 137.0407 | 10.3 |
| Asymmetric dimethylarginine (ADMA) | light | C8H18N4O2 | [M-H] | 201.1357 | 11.30 |
| ATP | light | C10H16N5O13P3 | [M-H] | 505.9885 | 11.3 |
| beta-alanine | light | C3H7NO2 | [M+H] | 90.0550 | 9.8 |
| betaine | light | C5H11NO2 | [M+H] | 118.0863 | 6.05 |
| Butyrylcarnitine | light | C11H21NO4 | [M+H] | 232.1543 | 3.7 |
| C5 Carnitines | light | C12H23NO4 | [M+H] | 246.1700 | 3.26 |
| cAMP | light | C10H12N5O6P | [M+H] | 330.0598 | 5.44 |
| Carbamoyl Aspartate | light | C5H8N2O5 | [M-H] | 175.0360 | 11.7 |
| carnitine | light | C7H15NO3 | [M+H] | 162.1125 | 8 |
| carnosine | light | C9H14N4O3 | [M+H] | 227.1139 | 9.8 |
| CDP | light | C9H15N3O11P2 | [M-H] | 402.0109 | 11.76 |
| CDP-choline | light | C14H26N4O11P2 | [M+H] | 489.1152 | 10.58 |
| choline | light | C5H14NO | [M+H] | 104.1070 | 10.1 |
| cis-aconitate | light | C6H6O6 | [M+H] | 175.0237 | 12.5 |
| cis-urocanic acid | light | C6H6N2O2 | [M+H] | 139.0502 | 4.01 |
| Citraconic acid | light | C5H6O4 | [M-H] | 129.0193 | 9.32 |
| citrate | light | C6H8O7 | [M-H] | 191.0197 | 12.5 |
| citrulline | light | C6H13N3O3 | [M+H] | 176.1030 | 10 |
| CMP | light | C9H14N3O8P | [M-H] | 322.0446 | 11.1 |
| Coenzyme A | light | C21H36N7O16P3S | [M+H] | 768.1225 | 9.5 |
| creatine | light | C4H9N3O2 | [M+H] | 132.0768 | 9.1 |
| creatinine | light | C4H7N3O | [M+H] | 114.0662 | 4.64 |
| CTP | light | C9H16N3O14P3 | [M-H] | 481.9772 | 12.6 |
| cystathionine | light | C7H14N2O4S | [M+H] | 223.0747 | 11.1 |
| cysteine | light | C3H7NO2S | [M-H] | 120.0125 | 4.8 |
| cystine | light | C6H12N2O4S2 | [M+H] | 241.0311 | 11 |
| cytidine | light | C9H13N3O5 | [M+H] | 244.0928 | 6.63 |
| cytosine | light | C4H5N3O | [M+H] | 112.0505 | 5.88 |
| dADP | light | C10H15N5O9P2 | [M+H] | 412.0418 | 10.05 |
| dAMP | light | C10H14N5O6P | [M+H] | 332.0754 | 9.01 |
| dATP | light | C10H16N5O12P3 | [M+H] | 492.0081 | 10.80 |
| dCDP | light | C9H15N3O10P2 | [M+H] | 388.0305 | 11.21 |
| dCMP | light | C9H14N3O7P | [M+H] | 308.0642 | 10.24 |
| dCTP | light | C9H16N3O13P3 | [M-H] | 465.9823 | 11.8 |
| deoxyadenosine | light | C10H13N5O3 | [M+H] | 252.1091 | 3.70 |
| deoxycytidine | light | C9H13N3O4 | [M+H] | 228.0979 | 5.50 |
| deoxyguanosine | light | C10H13N5O4 | [M+H] | 268.1040 | 6.10 |
| deoxyuridine | light | C9H12N2O5 | [M-H] | 227.0673 | 4.40 |
| dGDP | light | C10H15N5O10P2 | [M+H] | 428.0367 | 11.65 |
| dGMP | light | C10H14N5O7P | [M+H] | 348.0704 | 11.14 |
| dGTP | light | C10H16N5O13P3 | [M-H] | 505.9885 | 12.37 |
| dihydrofolate | light | C19H21N7O6 | [M-H] | 442.1481 | 11.14 |
| dihydroorotate | light | C5H6N2O4 | [M-H] | 157.0255 | 6.7 |
| dihydroxyacetone_phosphate | light | C3H7O6P | [M-H] | 168.9908 | 10.8 |
| dTDP | light | C10H16N2O11P2 | [M-H] | 401.0157 | 10.14 |
| dTMP | light | C10H15N2O8P | [M-H] | 321.0493 | 9 |
| dTTP | light | C10H17N2O14P3 | [M-H] | 480.9820 | 11.4 |
| dUDP | light | C9H14N2O11P2 | [M-H] | 387.0000 | 11.00 |
| dUMP | light | C9H13N2O8P | [M-H] | 307.0337 | 9.64 |
| dUTP | light | C9H15N2O14P3 | [M-H] | 466.9663 | 11.47 |
| erythrose_4-phosphate | light | C4H9O7P | [M-H] | 199.0013 | 11.35 |
| F6P/G1P | light | C6H13O9P | [M-H] | 259.0224 | 11.2 |
| FAD | light | C27H33N9O15P2 | [M-H] | 784.1499 | 7.78 |
| folate | light | C19H19N7O6 | [M-H] | 440.1324 | 12.16 |
| Formiminoglutamic acid (FIGLU) | light | C6H10N2O4 | [M+H] | 175.0713 | 9.88 |
| Fructose 1,6-bisphosphate | light | C6H14O12P2 | [M+H] | 341.0033 | 12.87 |
| fumarate | light | C4H4O4 | [M-H] | 115.0037 | 10.22 |
| gamma-aminobutyric acid (GABA) | light | C4H9NO2 | [M+H] | 104.0706 | 9.88 |
| GAR | light | C7H15N2O8P | [M-H] | 285.0493 | 11.55 |
| g-Butyrobetaine | light | C7H15NO2 | [M+H] | 146.1175 | 8.14 |
| GDP | light | C10H15N5O11P2 | [M-H] | 442.0170 | 12.7 |
| D-Glucose | light | C6H12O6 | [M-H] | 179.0561 | 9.45 |
| Glucose 6-phosphate | light | C6H13O9P | [M-H] | 259.0224 | 11.88 |
| Glucuronic acid | light | C6H10O7 | [M-H] | 193.0354 | 11.23 |
| glutamate | light | C5H9NO4 | [M-H] | 146.0459 | 10.3 |
| glutamate-13C5,15N | heavy | C'5H9N'O4 | [M-H] | 152.0597 | 10.3 |
| glutamine | light | C5H10N2O3 | [M+H] | 147.0764 | 9.60 |
| glutamyl-alanine | light | C8H14N2O5 | [M+H] | 219.0975 | 10 |
| glyceraldehyde_3-phosphate | light | C3H7O6P | [M-H] | 168.9908 | 11.35 |
| Glycerol 3-Phosphate | light | C3H9O6P | [M+H] | 173.0210 | 10.53 |
| Glycerophosphocholine | light | C8H20NO6P | [M+H] | 258.1101 | 9.53 |
| glycerophosphoethanolamine | light | C5H14NO6P | [M+H] | 216.0632 | 10.07 |
| glycine | light | C2H5NO2 | [M+H] | 76.0393 | 10.00 |
| glycine_13C2-15N | heavy | C'2H5N'O2 | [M+H] | 79.0430 | 10.00 |
| GMP | light | C10H14N5O8P | [M-H] | 362.0507 | 11.80 |
| GPI/glycerophosphoinositol | light | C9H19O11P | [M+H] | 335.0738 | 11.2 |
| GSH | light | C10H17N3O6S | [M+H] | 308.0911 | 10.12 |
| GSSG | light | C20H32N6O12S2 | [M+H] | 613.1592 | 12.28 |
| GTP | light | C10H16N5O14P3 | [M-H] | 521.9834 | 13.23 |
| guanidinoacetate | light | C3H7N3O2 | [M+H] | 118.0611 | 9.88 |
| guanine | light | C5H5N5O | [M+H] | 152.0567 | 6.9 |
| guanosine | light | C10H13N5O5 | [M+H] | 284.0989 | 7.33 |
| histidine | light | C6H9N3O2 | [M+H] | 156.0768 | 9.12 |
| histidine-13C6-15N3 | heavy | C'6H9N'3O2 | [M+H] | 165.0880 | 9.12 |
| homocysteine | light | C4H9NO2S | [M+H] | 136.0427 | 8.00 |
| hydroxyproline | light | C5H9NO3 | [M+H] | 132.0655 | 9.27 |
| hypotaurine | light | C2H7NO2S | [M+H] | 110.0270 | 9.47 |
| hypoxanthine | light | C5H4N4O | [M+H] | 137.0458 | 5.2 |
| IMP | light | C10H13N4O8P | [M-H] | 347.0398 | 10.64 |
| inosine | light | C10H12N4O5 | [M+H] | 269.0881 | 6.11 |
| isoleucine | light | C6H13NO2 | [M+H] | 132.1019 | 6.05 |
| Isoleucine-13C6-15N | heavy | C'6H13N'O2 | [M+H] | 139.1191 | 6.05 |
| Itaconic Acid | light | C5H6O4 | [M-H] | 129.0193 | 10.77 |
| Kynurenic Acid | light | C10H7NO3 | [M+H] | 190.0499 | 3.2 |
| kynurenine | light | C10H12N2O3 | [M+H] | 209.0921 | 5.6 |
| lactate | light | C3H6O3 | [M-H] | 89.0244 | 5.56 |
| leucine | light | C6H13NO2 | [M+H] | 132.1019 | 5.64 |
| leucine-13C6-15N | heavy | C'6H13N'O2 | [M+H] | 139.1191 | 5.64 |
| lysine | light | C6H14N2O2 | [M+H] | 147.1128 | 13.59 |
| lysine-13C6-15N2 | heavy | C'6H14N'2O2 | [M+H] | 155.1270 | 13.59 |
| malate | light | C4H6O5 | [M-H] | 133.0143 | 11.24 |
| methionine | light | C5H11NO2S | [M+H] | 150.0583 | 6.39 |
| Methionine sulfoxide | light | C5H11NO3S | [M+H] | 166.0532 | 8.25 |
| methionine-13C5-15N | heavy | C'5H11N'O2S | [M+H] | 156.0721 | 6.39 |
| methylmalonylcarnitine | light | C11H19NO6 | [M+H] | 262.1285 | 8.2 |
| methylthioadenosine | light | C11H15N5O3S | [M+H] | 298.0968 | 3.40 |
| myo-Inositol | light | C6H12O6 | [M-H] | 179.0561 | 11.20 |
| Na_acetylglutamine | light | C7H12N2O4 | [M+H] | 189.0870 | 6.60 |
| Na-acetylornithine | light | C7H14N2O3 | [M+H] | 175.1077 | 9.65 |
| N-acetylcysteine | light | C5H9NO3S | [M-H] | 162.0230 | 4.30 |
| N-acetylglutamate | light | C7H11NO5 | [M+H] | 190.0710 | 9.93 |
| N-Acetyl-L-aspartic acid | light | C6H9NO5 | [M+H] | 176.0554 | 10.34 |
| N-acetylmethionine | light | C7H13NO3S | [M+H] | 192.0689 | 3.00 |
| N-acetylphenylalanine | light | C11H13NO3 | [M+H] | 208.0968 | 2.34 |
| NAD+ | light | C21H28N7O14P2 | [M+H] | 664.1164 | 9.68 |
| NADH | light | C21H29N7O14P2 | [M-H] | 664.1175 | 9.36 |
| NADP | light | C21H29N7O17P3 | [M+H] | 744.0827 | 11.92 |
| NADPH | light | C21H30N7O17P3 | [M+H] | 746.0984 | 12.25 |
| Ne-acetyllysine | light | C8H16N2O3 | [M+H] | 189.1234 | 7.81 |
| nicotinamide | light | C6H6N2O | [M+H] | 123.0553 | 3.20 |
| Nicotinic Acid | light | C6H5NO2 | [M+H] | 124.0393 | 4.20 |
| O-adipoylcarnitine | light | C13H23NO6 | [M+H] | 290.1598 | 8.3 |
| ornithine | light | C5H12N2O2 | [M+H] | 133.0972 | 12.70 |
| orotate | light | C5H4N2O4 | [M-H] | 155.0098 | 6.14 |
| O-succinylcarnitine | light | C11H19NO6 | [M+H] | 262.1285 | 8.8 |
| palmitate | light | C16H32O2 | [M-H] | 255.2330 | 2.05 |
| p-Aminobenzoate | light | C7H7NO2 | [M-H] | 136.0404 | 6.80 |
| Pantothenic Acid | light | C9H17NO5 | [M+H] | 220.1180 | 4.50 |
| Phenylacetyl glutamine | light | C13H16N2O4 | [M+H] | 265.1183 | 3.40 |
| phenylalanine | light | C9H11NO2 | [M+H] | 166.0863 | 5.05 |
| phenylalanine-13C9-15N | heavy | C'9H11N'O2 | [M+H] | 176.1135 | 5.05 |
| phosphocholine | light | C5H14NO4P | [M+H] | 184.0733 | 10.17 |
| phosphocreatine | light | C4H10N3O5P | [M+H] | 212.0431 | 10.75 |
| phosphoenolpyruvate (PEP) | light | C3H5O6P | [M-H] | 166.9751 | 12.40 |
| phosphoethanolamine | light | C2H8NO4P | [M+H] | 142.0264 | 11.00 |
| phosphoserine | light | C3H8NO6P | [M-H] | 184.0017 | 11.60 |
| Pipecolic Acid | light | C6H11NO2 | [M+H] | 130.0863 | 7.10 |
| proline | light | C5H9NO2 | [M+H] | 116.0706 | 7.53 |
| proline-13C5-15N | heavy | C'5H9N'O2 | [M+H] | 122.0844 | 7.53 |
| propionylcarnitine | light | C10H19NO4 | [M+H] | 218.1387 | 5.26 |
| PRPP | light | C5H13O14P3 | [M-H] | 388.9445 | 13.21 |
| pseudouridine | light | C9H12N2O6 | [M-H] | 243.0623 | 6.91 |
| Pyridoxal | light | C8H9NO3 | [M+H] | 168.0655 | 4.2 |
| Pyridoxine | light | C8H11NO3 | [M+H] | 170.0812 | 3.8 |
| pyruvate | light | C3H4O3 | [M-H] | 87.0088 | 4.45 |
| Ribose 1-phosphate | light | C5H11O8P | [M-H] | 229.0119 | 10.80 |
| Ribose/Ribulose 5-phosphate | light | C5H11O8P | [M-H] | 229.0119 | 11.18 |
| saccharopine | light | C11H20N2O6 | [M+H] | 277.1394 | 10.99 |
| S-Adenosyl-L-Homocysteine (SAH) | light | C14H20N6O5S | [M+H] | 385.1289 | 8.44 |
| S-Adenosyl-L-methionine (SAM) | light | C15H22N6O5S | [M+H] | 399.1445 | 9.94 |
| sarcosine | light | C3H7NO2 | [M+H] | 90.0550 | 8.57 |
| Sedoheptulose-7-phosphate (D) | light | C7H15O10P | [M-H] | 289.0330 | 11.42 |
| serine | light | C3H7NO3 | [M+H] | 106.0499 | 10.00 |
| serine-13C3-15N | heavy | C'3H7N'O3 | [M+H] | 110.0570 | 10.00 |
| S-methylcysteine | light | C4H9NO2S | [M+H] | 136.0427 | 7.20 |
| sorbitol | light | C6H14O6 | [M-H] | 181.0718 | 8.77 |
| succinate | light | C4H6O4 | [M-H] | 117.0193 | 10.59 |
| taurine | light | C2H7NO3S | [M-H] | 124.0074 | 9.61 |
| threonine | light | C4H9NO3 | [M+H] | 120.0655 | 9.18 |
| threonine-13C4-15N | heavy | C'4H9N'O3 | [M+H] | 125.0760 | 9.18 |
| thymidine | light | C10H14N2O5 | [M-H] | 241.0830 | 3.20 |
| thymine | light | C5H6N2O2 | [M+H] | 127.0502 | 3.45 |
| TMAO | light | C3H9NO | [M+H] | 76.0757 | 5.85 |
| trans-urocanic acid | light | C6H6N2O2 | [M+H] | 139.0502 | 6.74 |
| Trimethyllysine | light | C9H20N2O2 | [M+H] | 189.1598 | 12.1 |
| tryptophan | light | C11H12N2O2 | [M+H] | 205.0972 | 6.31 |
| tyrosine | light | C<sub>9</sub>H<sub>11</sub>NO<sub>3</sub> | [M+H] | 182.0812 | 7.83 |
| tyrosine-13C<sub>9</sub>-15N | heavy | C'<sub>9</sub>H<sub>11</sub>N'O<sub>3</sub> | [M+H] | 192.1084 | 7.83 |
| UDP | light | C<sub>9</sub>H<sub>14</sub>N<sub>2</sub>O<sub>12</sub>P<sub>2</sub> | [M-H] | 402.9949 | 11.35 |
| UDP-GlcNAc | light | C<sub>17</sub>H<sub>27</sub>N<sub>3</sub>O<sub>17</sub>P<sub>2</sub> | [M+H] | 608.0889 | 10.81 |
| UDP-hexose | light | C<sub>15</sub>H<sub>24</sub>N<sub>2</sub>O<sub>17</sub>P<sub>2</sub> | [M-H] | 565.0477 | 11.74 |
| UMP | light | C<sub>9</sub>H<sub>13</sub>N<sub>2</sub>O<sub>9</sub>P | [M-H] | 323.0286 | 10.75 |
| uracil | light | C<sub>4</sub>H<sub>4</sub>N<sub>2</sub>O<sub>2</sub> | [M-H] | 111.0200 | 4.00 |
| urate | light | C<sub>5</sub>H<sub>4</sub>N<sub>4</sub>O<sub>3</sub> | [M-H] | 167.0211 | 8.60 |
| uridine | light | C<sub>9</sub>H<sub>12</sub>N<sub>2</sub>O<sub>6</sub> | [M-H] | 243.0623 | 5.25 |
| UTP | light | C<sub>9</sub>H<sub>15</sub>N<sub>2</sub>O<sub>15</sub>P<sub>3</sub> | [M-H] | 482.9613 | 12.50 |
| valine | light | C<sub>5</sub>H<sub>11</sub>NO<sub>2</sub> | [M+H] | 118.0863 | 7.15 |
| valine-13C<sub>5</sub>-15N | heavy | C'<sub>5</sub>H<sub>11</sub>N'O<sub>2</sub> | [M+H] | 124.1001 | 7.15 |
| xanthine | light | C<sub>5</sub>H<sub>4</sub>N<sub>4</sub>O<sub>2</sub> | [M-H] | 151.0262 | 7.27 |
| xanthosine | light | C<sub>10</sub>H<sub>12</sub>N<sub>4</sub>O<sub>6</sub> | [M-H] | 283.0684 | 7.80 |
| XMP | light | C<sub>10</sub>H<sub>13</sub>N<sub>4</sub>O<sub>9</sub>P | [M-H] | 363.0191 | 12.42 |

